# Multi-omics profiling reveals ethylene signalling as a key pathway underlying both genetic and epigenetic responses to low-dose ionizing radiation in *Arabidopsis*

**DOI:** 10.1101/2023.10.14.562363

**Authors:** Jeremy R B Newman, Mingqi Zhou, J. Chris Pires, Rhonda L Bacher, Michael Kladde, Patrick Concannon

## Abstract

There is increasing interest in the effects of low-dose ionizing radiation (IR) on plants as might occur during spaceflight, or as a consequence of human activities, such as nuclear power generation, that may result in the release of radioactive materials into the environment. High IR doses have long been used for the induction of mutations in plants with the goal of generating desirable traits for agribusiness. Less is known about the responses of plants to acute low doses of IR exposure. Here, we take a multi-omics approach to characterize the response to low dose IR in *Arabidopsis thaliana*. We adapt the Methyltransferase Accessibility Protocol for individual templates (MAPit) technique for use in plants allowing us to assay the epigenetic response to acute low-dose IR (10 cGy and 100 cGy) 72 hr after exposure, and, in parallel, use RNA sequencing to profile the transcription response at 1, 3, 24 and 72 hr after exposure. IR exposures as low as 10 cGy elicit robust genetic responses in *A. thaliana* detectable as early as 1 hr after exposure. Further examination revealed dose-dependent changes in gene expression, chromatin accessibility and DNA methylation that implicate the ethylene signalling pathway and abiotic stress response as underlying the transcriptional and epigenetic changes associated with IR. These changes are observable up to 72 hr post-exposure, suggesting that they are maintained well after the initial acute exposure. Our findings indicate that *A. thaliana* executes a multi-modal response to low-dose IR through induction and regulation of the ethylene response pathway.

## Introduction

Ionizing radiation (IR) poses an existential threat for most living organisms due to its genotoxic effects. Specifically, IR generates DNA double-strand breaks (DSBs) that can compromise both the content and the structural integrity of DNA. In addition to directly damaging DNA, IR can cause indirect damage to DNA, protein and lipids through the production of reactive oxygen species (ROS) (Azzam et al. 2012). Levels of background radiation due to radionuclides in the continental crust were much higher than present day when life first appeared (Karam and Leslie 1999). To counter this threat, living organisms have evolved a variety of mechanisms that sense, signal the presence of, and respond to DNA DSBs. In eukaryotes, these responses include cell cycle arrest, DNA repair, and/or activation of apoptotic pathways.

Plants, such as *Arabidopsis thaliana*, have limited options for evading environmental stresses because of their sessile lifestyle. Like many other eukaryotes, in response to DNA DSBs induced by IR, *A. thaliana* mounts a rapid response mediated, in large part, by the serine-threonine kinases ATM (Ataxia Telangiectasia Mutated) and ATR (Ataxia Telangiectasia and Rad3 Related). These kinases localize to DSBs, phosphorylating a number of protein substrates and thereby activating multiple downstream response pathways (Culligan et al. 2006; Kim et al. 2019). However, the response of plants to IR is much broader than simply the immediate response to DNA damage and includes biochemical, physiological, and transcriptional responses (Gudkov et al. 2019). Several genes encoding epigenetic regulators of gene expression in *A. thaliana* also display increased expression after IR exposure, suggesting that plants may also mount epigenetic responses to IR exposure (Sidler et al. 2015). For example, in 14-day-old irradiated *A. thaliana* plants, Wang et al. (Wang et al. 2016) observed IR- and ATM-dependent increases in the expression of transposable elements (TEs) and long non-coding RNAs whose expression is regulated epigenetically.

There is a long history of the application of IR to plants for the induction of mutations with the goal of generating more desirable traits for agribusiness as well as in experimental settings to better understand IR effects on plants (Kim et al. 2019; Lal et al. 2020; Ma et al. 2021). These approaches have frequently utilized doses of IR in excess of 1 Gy. As a result, much of the controlled experimental data available regarding responses to IR in plants corresponds to such doses. Observational studies at sites with high natural or induced levels of radiation provide much of the available data regarding plant responses to lower doses of IR. There is increasing interest in the effects of lower doses of IR on plants as might occur during spaceflight, or as a consequence of human activities that result in the release of radioactive materials into the environment (Mousseau and Moller 2020; De Pascale et al. 2021).

In the current study, we characterized the transcriptional and epigenetic responses of leaves from 4-week-old *A. thaliana* plants at various time points after acute exposure to low doses of IR (10 cGy and 100 cGy). We adapted the Methyltransferase Accessibility Protocol for individual templates (MAPit) (Pardo et al. 2015), a single-molecule methylation footprinting assay, applying it for the first time to plants. MAPit allowed us to simultaneously measure changes in both chromatin accessibility and endogenous DNA methylation in response to IR exposure. RNA sequencing (RNA-seq) was performed in parallel to quantify gene expression. Integrating these datasets, we find that exposure of *A. thaliana* to as little as 10 cGy of IR results in a robust response detectable from 1 hr to 72 hr post-exposure. This response includes significant and coordinated changes in gene expression, chromatin accessibility, and DNA methylation. The responses are dose-dependent, both in terms of the genes or sites affected at a given dose and the trajectory of response for a given gene or dose over time. Based on differentials in both gene expression and accessibility of chromatin at binding site motifs for specific transcription factors (TFs), the response to low-dose IR is significantly enriched for genes involved in ethylene signalling and response to various abiotic stresses.

## Results

We sought to characterize the genetic response of *A. thaliana* to low doses of IR. We profiled the transcriptional response using RNA-seq to 10 cGy and 100 cGy IR at several time points after exposure (1 hr, 3 hr, 24 hr, and 72 hr) as well as the epigenetic response at 72 hr post-IR exposure using the MAPit protocol (Pardo et al. 2015) to assay endogenous methylation and chromatin accessibility (**Fig. 1A**). A noticeable phenotype that appeared 72 hr after exposure to IR was leaf chlorosis, specifically at the higher IR dose (**Fig. 1B**).

**Figure 1.**
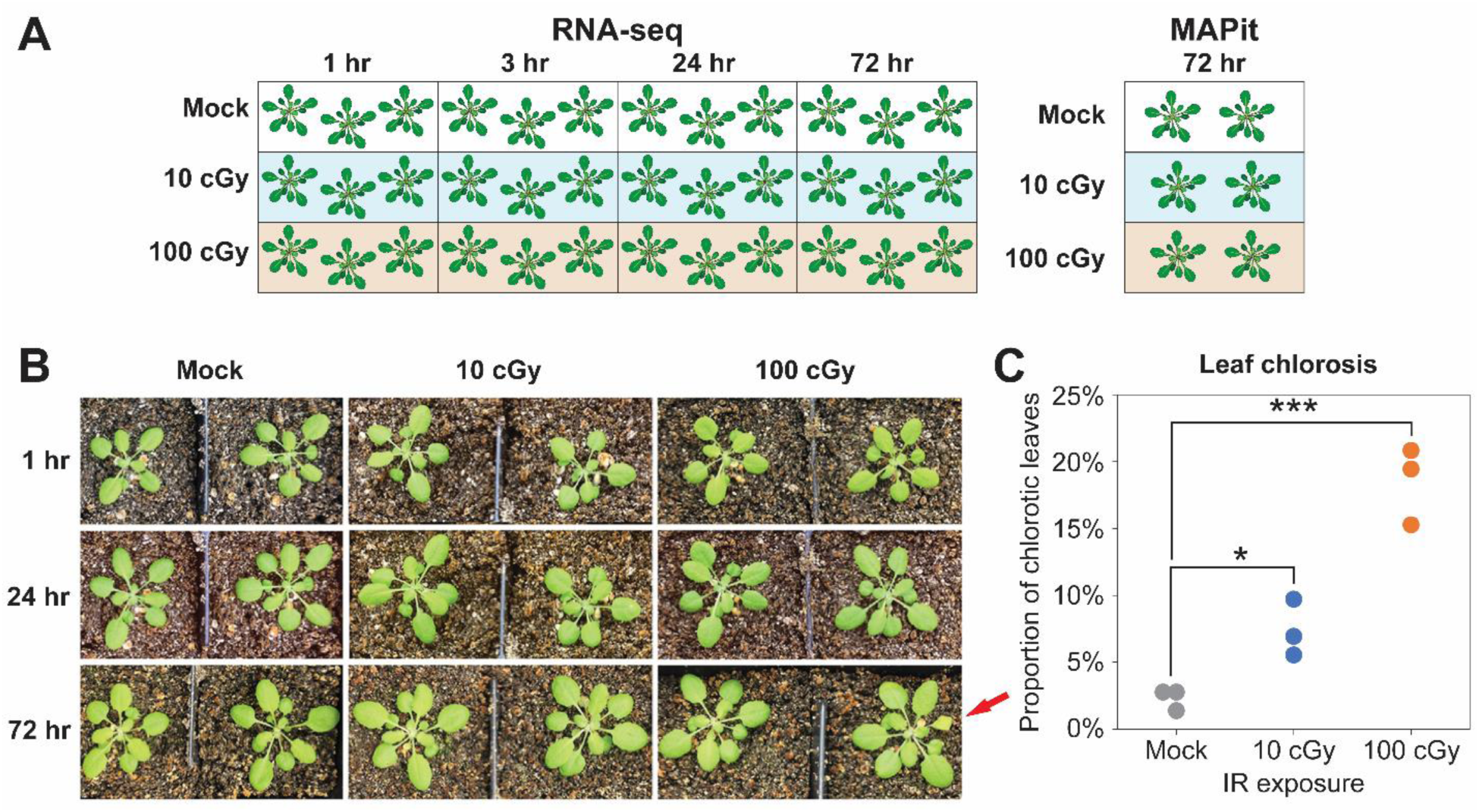
Experimental design. (**A**) Experiment design of RNA-seq and MAPit datasets showing dosimetry (Mock, 10 cGy, 100 cGy), time harvested after ionizing radiation (IR) exposure (1 hr, 3 hr, 24 hr, 72 hr), and number of replicates used. (**B**) *A. thaliana* plants after 1 hr, 24 hr, and 72 hr ionizing radiation exposure by dose. Red arrow highlights chlorosis at 72 hr post-exposure to 100 cGy of ionizing radiation. (C) The rate of leaf chlorosis in plants at 72 hr post-exposure to Mock (mean = 2.3 ± 0.8%), 10 cGy (mean = 7.4 ± 2.1%) and 100 cGy IR (mean = 18.5 ± 2.9%). Student’s *t*-test, * *P* = 0.02, *** *P* = 7.3×10^-4^).

### Exposure of *A. thaliana* to low-dose IR elicits dose-dependent transcriptional responses with diverging temporal trajectories

In order to identify genes expressed in *A. thaliana* whose expression was modulated following exposure to IR doses ≤ 100 cGy, RNA was extracted and sequenced from leaves of plants exposed to 10 cGy and 100 cGy of IR. A robust transcriptional response in terms of differentially expressed genes (DEGs) (log_2_ fold change ≥ 1 or ≤ −1 and FDR-corrected *P* < 0.05) was observed 1 hr following exposure to 10 cGy (489 DEGs of 22,889 expressed genes) or 100 cGy IR (343 DEGs of 22,889 expressed genes) (**Fig. 2A and B**). Across both IR exposures examined, at 1 hr after exposure, 449 genes were upregulated (log_2_ fold change ≥ 1) and 142 genes were downregulated (log_2_ fold change ≤ −1; **Fig. 2C**). Analysis of Gene Ontology annotations reveals that genes involved in ethylene signalling and hormone response peptide transport were overrepresented among genes that were upregulated at 1 hr, regardless of the dose (**Table 1**). Hierarchical clustering of all DEGs assigned 409 of the 449 DEGs upregulated at 1 hr in one of two clusters (**Fig. 2A)**. The first cluster contained 299 genes of which 288 genes were differentially expressed (DE) for either IR exposure at 1 hr. The expression of the genes in this cluster largely returned to baseline after 3 hr post-exposure. (**Fig. 2A**, Cluster 1). The second cluster comprised 145 genes of which 121 genes were DE at 1 hr (**Fig. 2A**, Cluster 2). The increase in expression 1 hr after IR exposure was overall larger than Cluster 1 and was observed to be generally maintained at the 3 hr timepoint returning to baseline by 24 hr (**Fig. 2A**). This second cluster of upregulated genes also had significantly more genes with a higher expression (log_2_ fold change ≥ 1, irrespective of *P* value) at 72 hr after IR exposure compared with the first cluster of DE genes (36% vs 1%; χ^2^ *P* = 2.48 × 10^-206^). A small number of additional DE genes (N = 45) were upregulated at 3 hr (**Fig. 2A**, Cluster 3) and were independent of those DE at 1 hr (**Fig. 2A**, Cluster 3). In contrast, most genes downregulated at 1 hr after exposure had their expression resolved by 3 hr. Six genes were consistently downregulated following IR exposure at all time points assayed – *FEP2*, *FEP3*, *AT2G14247*, *BHLH38*, *BHLH39*, and *BHLH100* – all of which encode proteins involved in iron homeostasis (Ashburner et al. 2000; Lamesch et al. 2012). Genes differentially expressed at 3 hr and at 24 hr were enriched for genes associated with iron homeostasis or terpenoid synthesis (**Table 1**).

**Figure 2.**
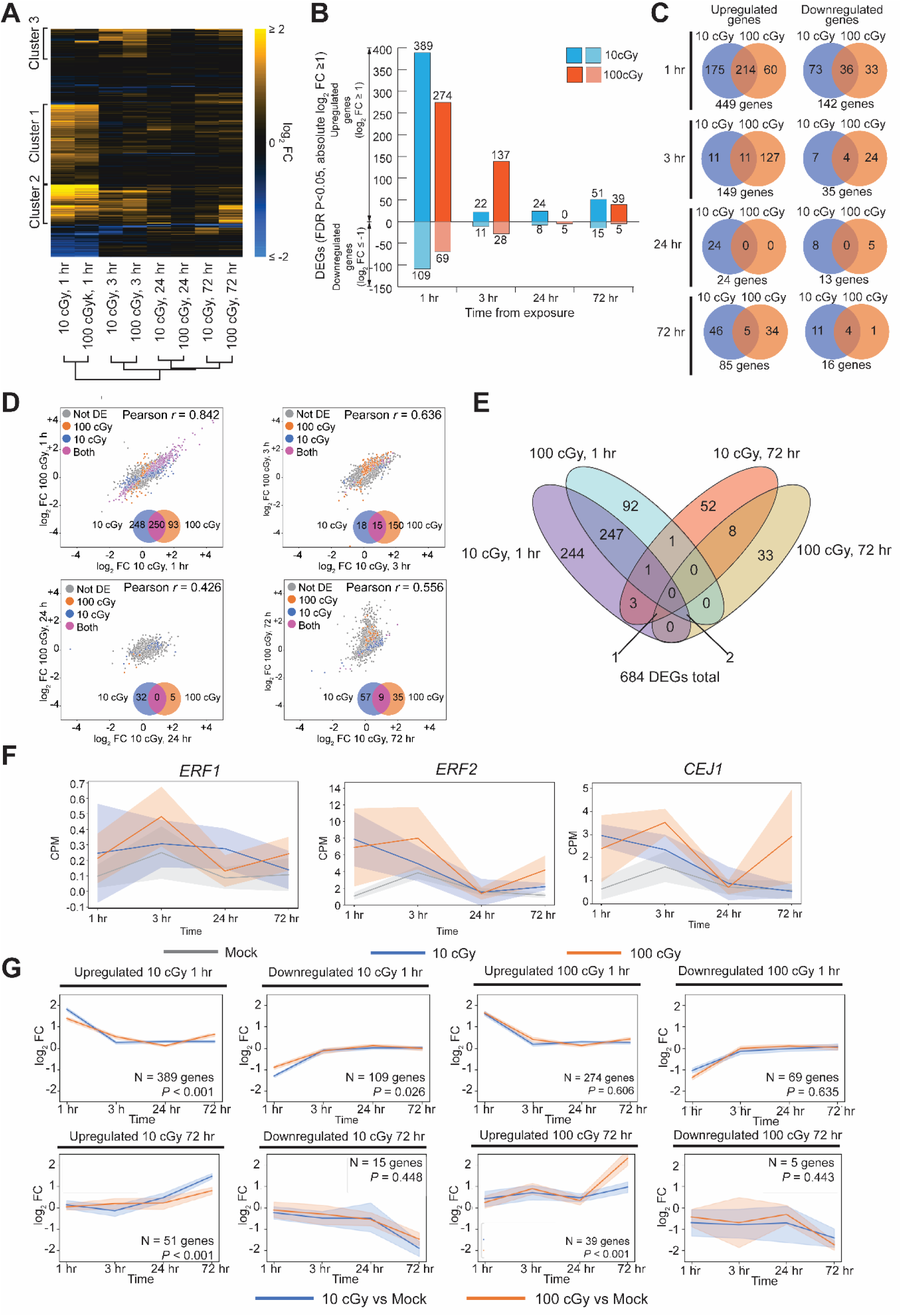
Transcriptional response to low-dose ionizing radiation. (**A**) Heatmap of differentially expressed genes (DEGs). (**B**) Up-regulated (log_2_ fold change (FC) ≥ 1 and FDR *P* < 0.05) and downregulated (log_2_ FC ≤ −1 and FDR *P* < 0.05) DEGs by timepoint. Blue bars represent the DEGs between 10 cGy and Mock, orange bars represent DEGs between 100 cGy and Mock. (**C**) Overlap between up- or downregulated DEGs between doses for each timepoint assayed. (**D**) Comparison of log_2_ FC between doses for each timepoint. Blue points are DEGs detected only at 10 cGy, orange points are DEGs only at 100 cGy, purple points are DEGs at both 10 cGy and 100 cGy, grey points are genes not differentially expressed at either dose (−1 < log_2_ FC < 1 and/or FDR *P* ≥ 0.05). (**E**) Overlap of DEGs at 1 hr and 72 hr at 10 cGy and 100 cGy. (**F**) Expression of the ethylene response factor genes *ERF1*, *ERF2*, and *CEJ1* over time. (**G**) Change in log_2_ FC over time for up- and downregulated DEGs at 1 h and 72 h for 10 cGy and 100 cGy. Blue lines represent log_2_ FC for 10 cGy vs Mock; orange lines represent log_2_ FC for 100 cGy vs Mock; shaded areas represent standard deviation.

**Table 1.**
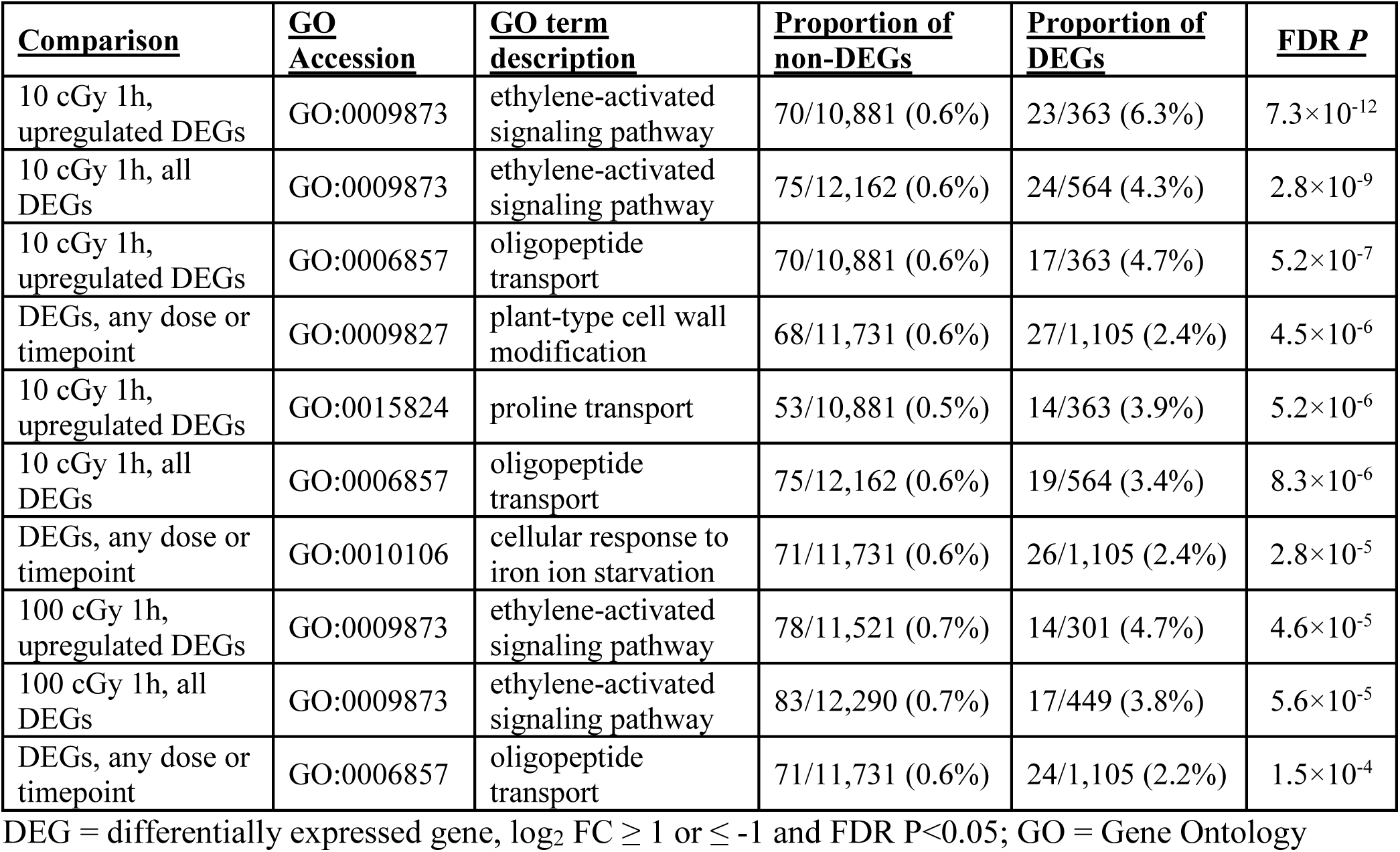
Top Gene Ontology enrichment of differentially expressed genes.

| <u>Comparison</u> | <u>GO Accession</u> | <u>GO term description</u> | <u>Proportion of non-DEGs</u> | <u>Proportion of DEGs</u> | <u>FDR *P*</u> |
| --- | --- | --- | --- | --- | --- |
| 10 cGy 1h, upregulated DEGs | GO:0009873 | ethylene-activated signaling pathway | 70/10,881 (0.6%) | 23/363 (6.3%) | $7.3 \times 10^{-12}$ |
| 10 cGy 1h, all DEGs | GO:0009873 | ethylene-activated signaling pathway | 75/12,162 (0.6%) | 24/564 (4.3%) | $2.8 \times 10^{-9}$ |
| 10 cGy 1h, upregulated DEGs | GO:0006857 | oligopeptide transport | 70/10,881 (0.6%) | 17/363 (4.7%) | $5.2 \times 10^{-7}$ |
| DEGs, any dose or timepoint | GO:0009827 | plant-type cell wall modification | 68/11,731 (0.6%) | 27/1,105 (2.4%) | $4.5 \times 10^{-6}$ |
| 10 cGy 1h, upregulated DEGs | GO:0015824 | proline transport | 53/10,881 (0.5%) | 14/363 (3.9%) | $5.2 \times 10^{-6}$ |
| 10 cGy 1h, all DEGs | GO:0006857 | oligopeptide transport | 75/12,162 (0.6%) | 19/564 (3.4%) | $8.3 \times 10^{-6}$ |
| DEGs, any dose or timepoint | GO:0010106 | cellular response to iron ion starvation | 71/11,731 (0.6%) | 26/1,105 (2.4%) | $2.8 \times 10^{-5}$ |
| 100 cGy 1h, upregulated DEGs | GO:0009873 | ethylene-activated signaling pathway | 78/11,521 (0.7%) | 14/301 (4.7%) | $4.6 \times 10^{-5}$ |
| 100 cGy 1h, all DEGs | GO:0009873 | ethylene-activated signaling pathway | 83/12,290 (0.7%) | 17/449 (3.8%) | $5.6 \times 10^{-5}$ |
| DEGs, any dose or timepoint | GO:0006857 | oligopeptide transport | 71/11,731 (0.6%) | 24/1,105 (2.2%) | $1.5 \times 10^{-4}$ |

DEG = differentially expressed gene, $\log_2$ FC $\geq 1$ or $\leq -1$ and FDR $P < 0.05$ ; GO = Gene Ontology

Changes in gene expression were similar for both IR doses with regard to the number of DEGs, the proportion of up- and downregulated genes (77% of DEGs upregulated at 10 cGy vs 81% of DEGs upregulated at 100 cGy; **Fig. 2B**) and fold change in expression (**Fig. 2D**). Of the 591 total DEGs at 1 hr, 42.3% (250 genes) were in common between the 10 cGy and 100 cGy doses (**Fig. 2D**). After this initial response, the transcriptional profiles at 10 cGy and 100 cGy diverged, with fewer of the total DEGs in common at each successive time point assayed (**Fig. 2C and D**). There were few DEGs after 10 cGy exposure at either 3 hr (33 DEGs) or 24 hr (32 DEGs) (**Fig. 2B**). At 100 cGy exposure, the number of DEGs at 3 hr (165 DEGs) was five times higher than for 10 cGy (33 DEGs) but returned to baseline expression by 24 hr post-exposure (5 DEGs) (**Fig. 2B**). Transcriptional differences between irradiated and mock-exposed plants re-emerged 72 hr after exposure, with a slightly higher response at the 10 cGy dose (**Fig. 2B and C**). Gene expression differences at 72 hr were mostly dose dependent as there were few genes in common between 10 cGy and 100 cGy (**Fig. 2C and D**). The DEGs at 72 hr were almost entirely distinct from those in the 1 hr response (**Fig. 2E**); however, several ethylene response factor genes that were upregulated at 1 hr (such as *ERF1*, *ERF2*, and *CEJ1*) were also upregulated at 72 hr (log_2_ fold change ≥ 1; **Fig. 2F**), although these genes did not reach statistical significance at this time point. Genes that were differentially expressed at 1 hr at either dose followed the same overall expression trajectory over time, whereas genes that were engaged later diverged in their expression by 72 hr (**Fig. 2G**), especially those that were upregulated at 72 hr at 100 cGy. No single biological process was highly enriched among the genes that respond to low-dose IR across or irrespective of time (**Supplemental Table 1**).

### Differences in chromatin accessibility are observable days after exposure to low-dose ionizing radiation

Gene expression changes indicated a common early response to low-dose IR that by 72 hr diverged into largely dose-dependent differences, suggesting that regulation and coordination of these responses should follow a similar trajectory. To examine regulatory changes after low-dose IR exposure we assayed the epigenetic response 72 hr after 10 cGy, 100 cGy, or mock exposure using the MAPit assay (Pardo et al. 2009; Pardo et al. 2015), which allows the simultaneous, genome-wide profiling of endogenous methylation and chromatin accessibility. With MAPit, chromatin accessibility is probed by treating purified nuclei with an exogenous DNA methyltransferase, M.CviPI (Xu et al. 1998 characterization and expression of the gene coding for a cytosine-5-DNA methyltransferase recognizing GpC), which recognizes GpC dinucleotides that can be modified by endogenous DNA methyltransferases in plants. Therefore, to derive differential chromatin accessibility in *A. thaliana*, we compared the difference between exogenous DNA methyltransferase-mediated and endogenous baseline methylated GC sites (Methods). **Supplemental Note 1** details our modification and optimization of the MAPit protocol for application to plants. A striking difference in chromatin accessibility, quantified as differentially accessible regions (DARs), was observed following exposure to IR. The response at the lower, 10 cGy dose was almost entirely comprised of hyperaccessible regions (hyperDARs, 99% at 10 cGy) compared to the response at 100 cGy (79%; **Fig. 3A**). The length of DARs detected at 10 cGy was similar to those detected at 100 cGy (10 cGy mean length = 170.7 ± 80.1 nt, median length = 161 nt; interquartile range = 114 – 214 nt; 100 cGy mean length = 161.1 ± 72.7 nt, median length = 154 nt; interquartile range = 107 – 208 nt; Kruskal-Wallis *P* = 0.002; **Fig. 3B**). Most DARs were dose-specific (**Fig. 3C**), though about 11% (425 of 3,920) of all significant hyperDARs were common to both IR exposures. Within hyperaccessible genic regions GC accessibility was ∼15% higher at 10 cGy exposure than at Mock or 100 cGy exposures. Toward the center of the genic DARs, the change in accessibility increased to 20-25% compared to Mock exposure; for 100 cGy this change was 15-20% (**Fig. 3D**). Compared to genic hyperDARs, the changes in GC accessibility in intergenic hyperDARs was less: ∼15% for 10 cGy and ∼10% for 100 cGy (**Fig. 3D**). This is consistent with genic regions being sites of greater chromatin remodeling activity. As there were few hypoaccessible regions (hypoDARs) at 10 cGy, our estimates of change in GC accessibility are necessarily less precise. However, on average, there was a 10% decrease in GC accessibility of hypoDARs for both 10 cGy and 100 cGy IR exposure (**Fig. 3E**); these average differences were greater at genic hypoDARs compare to intergenic hypoDARs.

**Figure 3.**
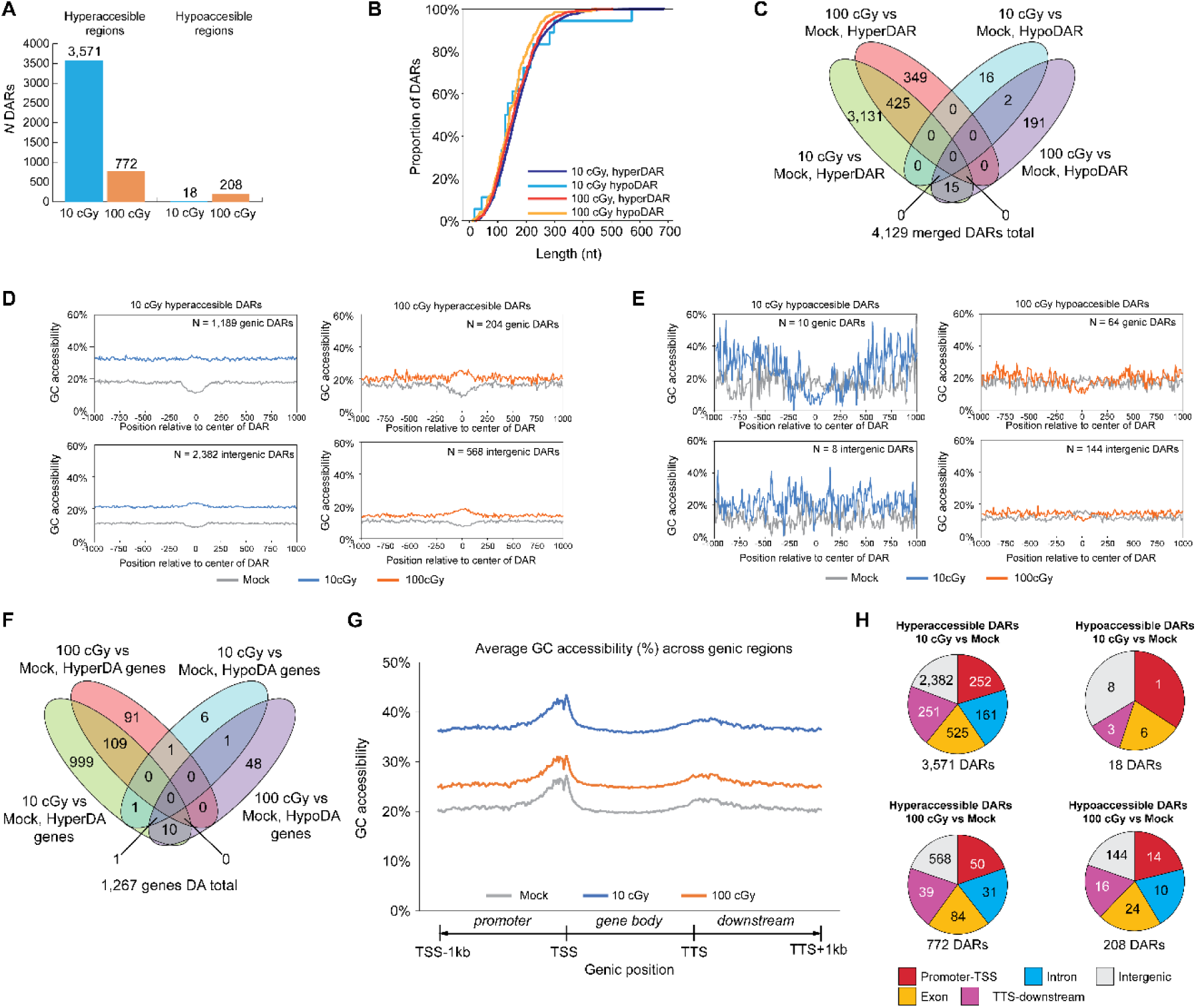
Differential chromatin accessibility in *A. thaliana* following low dose ionizing radiation exposure. (**A**) Number of significantly hyperaccessible regions (hyperDARs) and hypoaccessibile regions (hypoDARs) for 10 cGy and 100 cGy 72 hr after ionizing radiation (IR) exposure. (**B**) Distribution of differentially accessible region (DAR) length at 10 cGy and 100 cGy 72 hr after IR exposure. (**C**) Overlap of hyperDARs and hypoDARs for 10 cGy and 100 cGy 72 hr after IR exposure. (**D**) GC accessibility at genic (top) and intergenic (bottom) hyperDARs for 10 cGy and 100 cGy 72 hr after IR exposure. (**E**) GC accessibility at genic (top) and intergenic (bottom) hypoDARs for 10 cGy and 100 cGy 72 hr after IR exposure. (**F**) Overlap of hyper- and hypo-differentially accessible (DA) genes for 10 cGy and 100 cGy 72 hr after IR exposure. (**G**) Average GC accessibility across genic regions (1 kb 5’ promoter, gene body, 1 kb 3’ downstream). Length of gene bodies were rescaled to 1 kb. (**H**) Distribution of DARs by feature (promoter-transcriptional start site (TSS), exon, intron, transcriptional termination site (TTS)-downstream, intergenic). Proportion of pie chart is normalized to the total number of GC motifs within all DARs of that feature type.

When the set of genic DARs (1,199 DARs for 10 cGy, 268 DARs for 100 cGy) were considered based on their annotated genes, we found that like DARs, most differentially accessible (DA) genes were exposure-specific and effect-specific: 109 of the 1,212 (9%) of hyperaccessible genes and 2 of 68 (3%) of hypoaccessible genes were common to both IR doses (**Fig. 3F**). Few genes (13 of 1,267 DA genes (1%) displayed both regions of hyperaccessibility and regions of hypoaccessibility, demonstrating that most DA genes have a single regulatory fate. Examining average accessibility over genic regions (gene body ± 1 kb), a similar pattern at all exposures (10 cGy, 100 cGy, Mock) was observed (**Fig. 3G**). Despite both 10 cGy and 100 cGy IR exposures having a higher GC accessibility compared to Mock exposed, there is a common increase in GC accessibility corresponding to the sequence 5’ to the transcriptional start site (TSS) of a gene; this is consistent with the TSS being a site of active chromatin remodeling and regulation (**Fig. 3G**). While GC accessibility was otherwise generally consistent across the gene body, most DARs in genic regions were annotated to exons, then 3’ downstream and 5’ promoter regions (**Fig. 3H**). When normalized for the number of GC motifs in DARs annotated to each feature type, the distribution of annotated feature types to DARs was relatively consistent (**Fig. 3H**), suggesting that there is little bias in DAR detection for any one feature type.

A motif enrichment analysis of DARs revealed that DNA recognition sequences of several transcription factors were more frequently observed in hyperDARs after exposure to either 10 cGy or 100 cGy of IR. Consistent with the gene expression results, several of the highest scoring motifs included binding sites for transcription factors that are associated with ethylene signaling (ERF1, RRTF1, RAP2.3; **Fig. 4A** and **Supplemental Table 2**). Many of these transcription factor binding sites are also enriched in the promoter regions of DEGs (**Supplemental Table 3**). We then interrogated the RNA-seq data for genes associated with these transcription factor binding sites (**Supplemental Table 4**). We observed that many of these transcription factors are upregulated early (1 hr or 3 hr) following irradiation and several maintained elevated expression over time (**Fig. 4B**). We also examined the expression of several target genes for RAP2.3, RAP2.6, RRTF1, and ZAT18, that were DE for at least one dose and at least one time point examined. We found that the expression pattern of these target genes mirrored the expression of their associated transcription factors. Examples of these include *ATGILP* (RAP2.3, RAP2.6, RRTF1), *AT3G01830* (RAP2.3, RAP2.6, RRTF1), *SIP4* (ERF1, RAP2.3, RAP2.6, RRTF1), and *AT1G61380* (ZAT18, ZAT2) (**Fig. 4C**).

**Figure 4.**
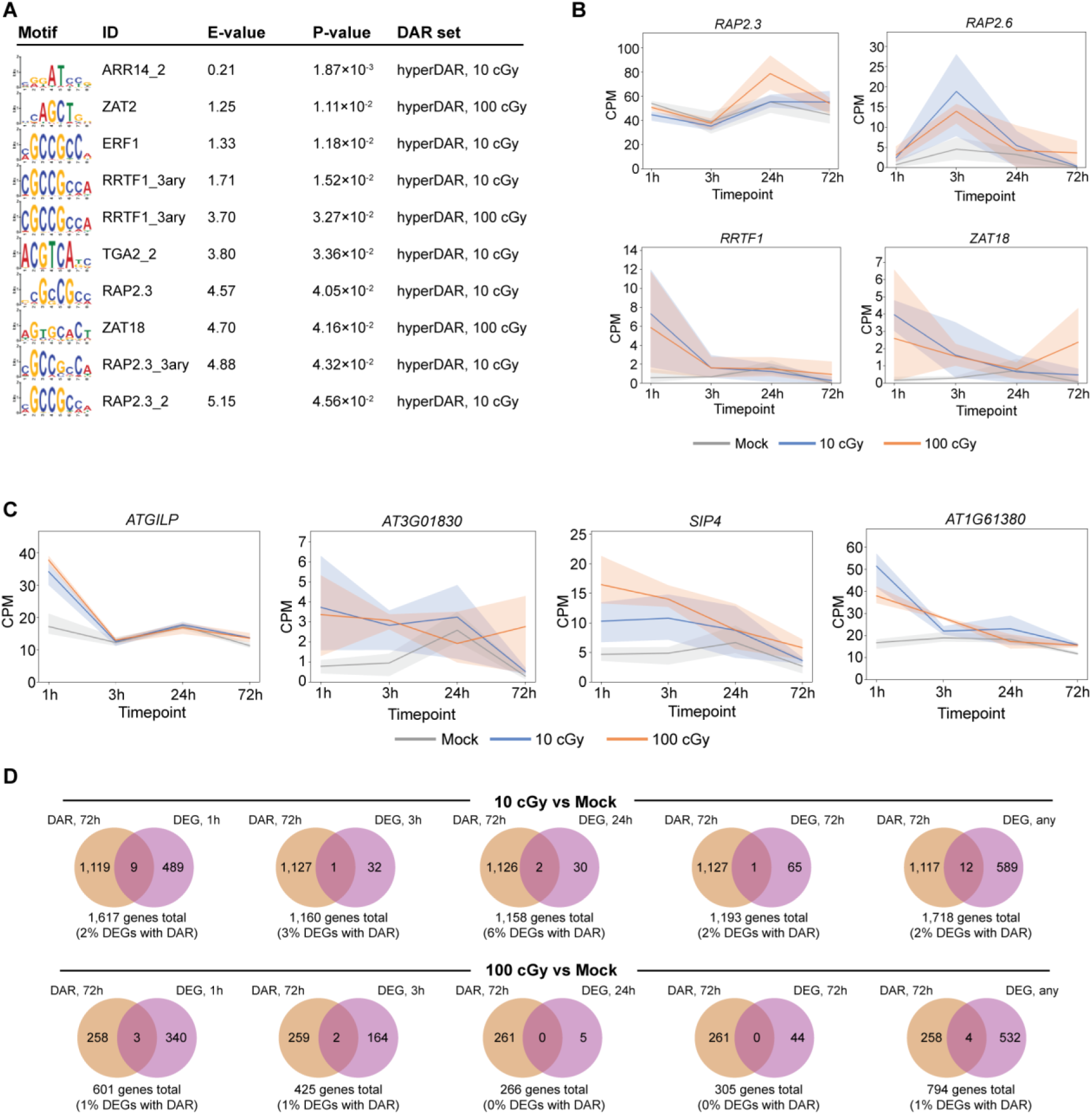
Comparison of differentially accessible regions and differentially expressed genes in Arabidopsis following ionizing radiation exposure. (**A**) Top 10 (by *P* value) motifs enriched in DARs (see **Supplemental Table 2** for full listing). (**B**) Expression of transcription factor genes *RRTF1*, *RAP2.3*, *RAP2.6* and *ZAT18* over 72 hr following exposure to ionizing radiation (IR). Shaded areas correspond to the 95% confidence interval of each gene’s expression. (**C**) Expression of *ATGLIP*, *AT3G01830*, *SIP4*, and *AT1G61380* over 72 hr following IR exposure. Shaded areas correspond to the 95% confidence interval of each gene’s expression. (**D**) Venn diagrams showing the overlap between differentially expressed genes (DEGs) and genes annotated to at least one DAR (distance between gene and DAR ±1 kb) for each IR dose (10 cGy, 100 cGy) and timepoint (1 hr, 3 hr, 24 hr, 72 hr, any timepoint assayed).

We also compared DARs at 72 hr and their most proximal (annotated) genes with the set of DEGs observed over the time series. Overall, few DEGs at 10 cGy or 100 cGy had at least one DAR within their annotated genic region (1 kb promoter, gene body, 1 kb downstream). These proportions were generally consistent between each time point (**Fig. 4D**).

### IR alters endogenous DNA methylation in *A. thaliana*

Our implementation of MAPit in plants allowed us to explore if the response to IR observed in transcription and chromatin accessibility also extended to other forms of epigenetic regulation. The expression of several DNA methyltransferases, such as *CMT2*, *CMT3*, and *MET1*, varied over the time course assayed after IR exposure, suggesting that we might expect to see changes in endogenous DNA methylation in response to IR exposure (**Supplemental Fig. 1**).

Indeed, robust changes in endogenous methylation in following exposure to IR were observed comparing samples not treated with M.CviPI: 71,687 DMRs were detected 72 hr after 10 cGy IR exposure, and 45,329 DMRs after 100 cGy IR exposure when compared to Mock-exposed plants (**Fig. 5A**). These methylation changes were largely at regions containing CHH motifs (81% of 10 cGy DMRs, 97% of 100 cGy DMRs; **Fig. 5A**). The changes in methylation were dose dependent in terms of both site type and direction of effect: CG and CHG methylation changes were more abundant at 10 cGy and more frequently corresponded to regions of hypomethylation (68% of 10 cGy CG DMRs hypomethylated, 88% of 10 cGy CHG DMRs hypomethylated, **Fig. 5B**). Changes in CG and CHG methylation at 100 cGy IR exposure were less extensive, with fewer total DMRs detected. For CHG DMRs, but not CG DMRs, there was less bias toward hypomethylation at 100 cGy than at the 10 cGy IR dose (78% of 100 cGy CG

**Figure 5.**
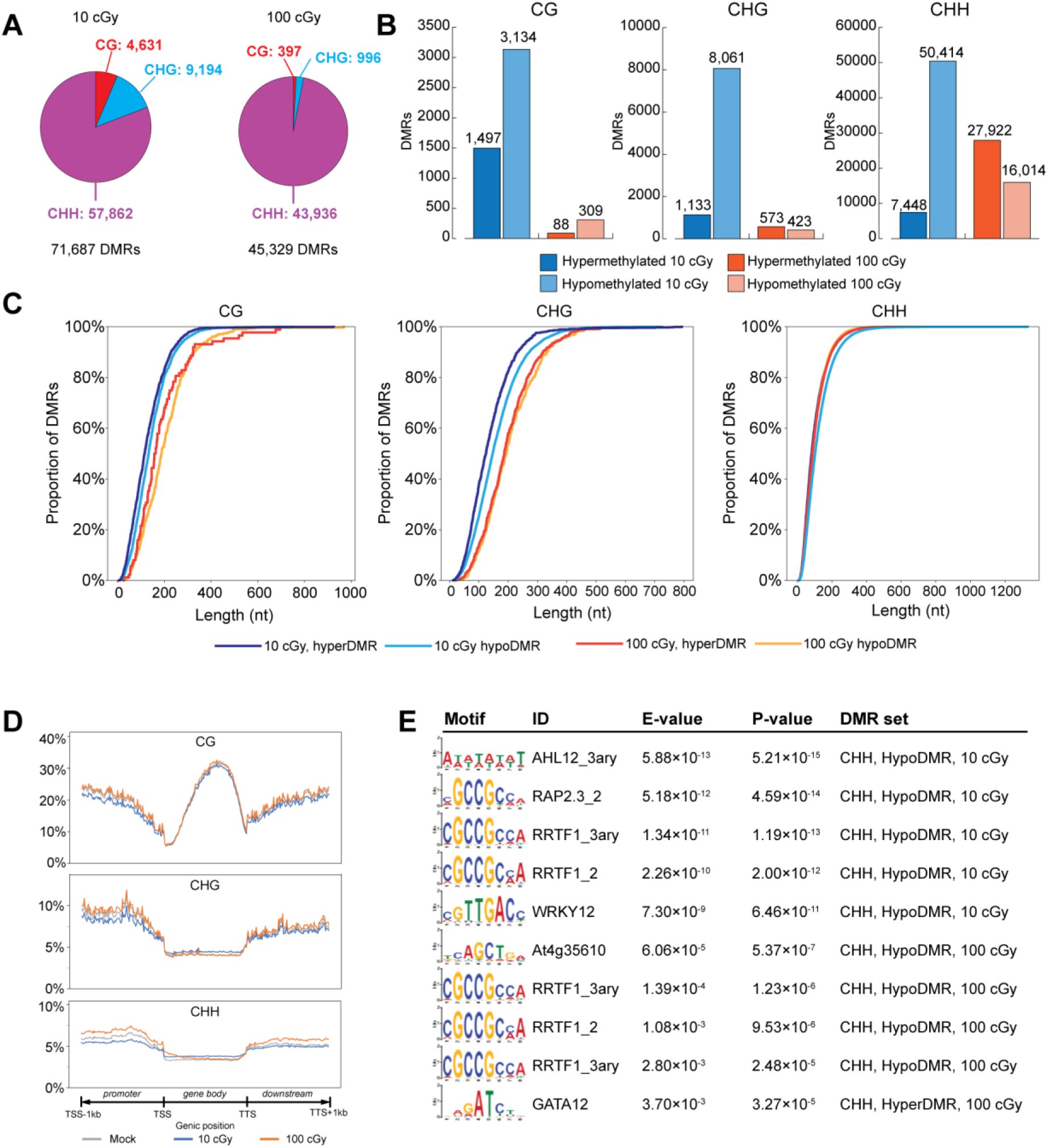
Differential endogenous methylation in *A. thaliana* following low dose ionizing radiation exposure. (**A**) Distribution of differentially methylated regions (DMRs) by site type and ionizing radiation (IR) exposure. (**B**) Distribution of hypermethylated and hypomethylated DMRs by site type and IR exposure. (**C**) Distribution of DMR length at 10 cGy and 100 cGy 72 hr after IR exposure. (**D**) Average CG, CHG, and CHH methylation across genic regions (1 kb 5’ promoter, gene body, 1 kb 3’ downstream). Gene body lengths were rescaled to 1 kb. (**E**) Top 10 (by *P* value) motifs enriched DMRs (see **Supplemental Table 5** for full table).

DMRs hypomethylated, 42% of 100 cGy CHG DMRs hypomethylated, **Fig. 5B**). Changes in CHH methylation followed a similar pattern as CHG methylation: the response at 10 cGy IR being larger and more biased toward hypomethylation compared to the response at 100 cGy IR (87% of 57,862 CHH DMRs hypomethylated at 10 cGy, 36% of 43,936 CHH DMRs hypomethylated at 100 cGy; χ^2^ *P* < 0.001; **Fig. 5B**). The average length of DMRs at 10 cGy tended to be shorter than those detected at 100 cGy (**Table 2**). While CG and CHG DMRs cover a similar range of lengths for both exposure, the average length for CG and CHG DMRs was significantly shorter at 10 cGy compared to 100 cGy (**Table 2**, **Fig. 5C**), indicating that methylation changes at the lower exposure tended to occur over smaller regions of the genome. CHH DMRs overall had a similar length distribution, except for 10 cGy hypoDMRs which were on average ∼20 nt longer than other CHH DMRs. The longest 10 cGy CHH DMR was also nearly twice the length of the longest 100 cGy CHH DMR, suggesting that changes in CHH methylation associated with IR exposure are different from that of CG or CHG methylation (**Table 2**, **Fig. 5**).

**Table 2.** Distribution of differentially methylated region lengths.

| IR exposure | Site type | N hyperDMRs / N hypoDMRs | DMR length range | Statistic | hyperDMR length (nt) | hypoDMR length (nt) | Kruskal-Wallis P |
| --- | --- | --- | --- | --- | --- | --- | --- |
| 10 cGy | CG | 1,497 / 3,134 | 6 to 925 nt | Mean (± SD) | 128.23 ± 79.61 | 143.33 ± 79.48 | $1.3 \times 10^{-40}$ |
|  |  |  |  | Median [Q<sub>1</sub> - Q<sub>3</sub>] | 155 [69 - 172] | 132 [86 - 184] |  |
| 100 cGy | CG | 88 / 309 | 25 to 969 nt | Mean (± SD) | 189.83 ± 124.73 | 202.56 ± 108.60 | |
|  |  |  |  | Median [Q<sub>1</sub> - Q<sub>3</sub>] | 160 [110.5 - 225.5] | 187 [125 - 254] |  |
| 10 cGy | CHG | 1,133 / 8,061 | 10 to 793 nt | Mean (± SD) | 136.93 ± 82.54 | 160.29 ± 85.42 | $4.3 \times 10^{-68}$ |
|  |  |  |  | Median [Q<sub>1</sub> - Q<sub>3</sub>] | 124 [80 - 177] | 147 [99 - 205] |  |
| 100 cGy | CHG | 573 / 423 | 21 to 718 nt | Mean (± SD) | 198.15 ± 95.15 | 205.39 ± 95.84 | |
|  |  |  |  | Median [Q<sub>1</sub> - Q<sub>3</sub>] | 187 [130 - 249] | 194 [137 - 261] |  |
| 10 cGy | CHH | 7,448 / 50,414 | 5 to 1,330 nt | Mean (± SD) | 101.34 ± 72.90 | 122.54 ± 83.83 | $< 1.0 \times 10^{-100}$ |
|  |  |  |  | Median [Q<sub>1</sub> - Q<sub>3</sub>] | 82 [50 - 132] | 102 [63 - 160] |  |
| 100 cGy | CHH | 27,922 / 16,014 | 6 to 759 nt | Mean (± SD) | 104.43 ± 70.67 | 103.37 ± 66.46 | |
|  |  |  |  | Median [Q<sub>1</sub> - Q<sub>3</sub>] | 87 [54 - 137] | 88 [55 - 135] |  |

IR = ionizing radiation; hyperDMR = hypermethylated region; hypoDMR = hypomethylated region; DMR = differentially methylated region

We collapsed and compared overlapping hyper- and hypoDMRs at 10 cGy and 100 cGy (**Supplemental Fig. 2**). Distinct separation of hyperDMRs and hypoDMRs was observed for CG and CHG regions. Forty percent of 100 cGy CG DMRs and 100 cGy CHG DMRs were shared with DMRs at 10 cGy, although these shared DMRs only made up about 4% of 10 cGy CG and CHG DMRs (**Supplemental Fig. 2**). When hyper- and hypomethylation is considered, a greater proportion of CHG hypoDMRs were shared between exposures (65% of 100 cGy CHG hypoDMRs) than were CHG hyperDMRs (18% of 100 cGy CHG hyperDMRs). Changes in CHH methylation were more extensive but less dose-dependent: 74% of 100 cGy hyperDMRs comprised 28% of 10 cGy hyperDMRs, and 76% of 100 cGy hypoDMRs comprised 25% of 10 cGy hypoDMRs. This indicates there is a common underlying epigenetic response, although there are more extensive dose-dependent effects at the lower exposures.

While more DMRs were located in intergenic regions than any single genic region feature (promoter-TSS, exon, intron, or TTS-downstream), when adjusted for the number of endogenously methylated sites, the proportion of differential methylation was relatively evenly distributed over different intergenic and genic features (**Supplemental Fig. 3**). Meta-analysis of all three types of the distribution of non-normalized endogenous methylation across genic regions showed characteristic minima in CG methylation at TSSs and TTSs, and CHG and CHH methylation were largely depleted from gene body regions (**Fig. 5D**). Qualitatively, similar patterns of endogenous methylation levels were seen over genic regions at different IR exposures (10 cGy, 100 cGy, and Mock). Quantitatively, on average, endogenous methylation, particularly CHG and CHH, at promoter regions and to a lesser extent 3’ downstream regions appeared to be lower at 10 cGy compared to Mock exposure (**Fig. 5D**, blue vs grey line), while promoter and downstream methylation was higher at 100 cGy compared to Mock (**Fig. 5D**, orange vs grey line). This dynamic is similar to the ratio of hyperaccessible to hypoaccessible regions seen at 10 cGy and 100 cGy (**Fig. 3A**), as well as the bias towards CHG and CHH hypomethylation at 10 cGy compared to 100 cGy (**Fig. 5B**).

We also performed a motif analysis on DMRs to ascertain what, if any, transcriptional networks they might regulate, as transcription factor binding can be sensitive to methylation status. Motif analysis on DMRs revealed that regions of CHH hypermethylation were enriched for DNA binding sequences of transcription factors associated with ethylene signaling (RRTF1, At4g35610, RAP2.3) or development/environmental responses (ARR14, AHL12. GATA12) (**Fig. 5E**; **Supplemental Table 5**). All of these factors, except for GATA12, were also in motif analysis of hyperDARs (**Supplemental Table 2**).

### Intersecting endogenous CG, CHG, and CHH methylation with chromatin accessibility reveals a common epigenetic response to IR

While the majority of observed DMRs were located in intergenic regions (**Supplemental Fig. 3**), in terms of genic features, the highest absolute fraction of CHH DMRs were located in promoter regions (∼50% of all genic CHH DMRs). This suggests the possibility that changes in CHH methylation may be regulating the expression of nearby genes, but relatively few DEGs had a DMR of any type within 1 kb (**Fig. 6A**). We then compared DARs and DMRs to determine what fraction of DARs overlapped with regions of altered methylation 72 hr after IR exposure. Relatively few DARs (1 - 7%) overlapped with CG DMRs after either exposure; 22% of 10 cGy DARs overlapped with at least one CHG DMR while only 4% of 100 cGy DARs overlapped with at least one CHG DMR at the same IR dose (**Fig. 6B**). At least half of DARs (50 - 59%) at both IR exposures overlapped with at least one CHH DMR (**Fig. 6B**), suggesting that changes in CHH methylation after IR exposure often accompany changes in chromatin accessibility.

**Figure 6.**
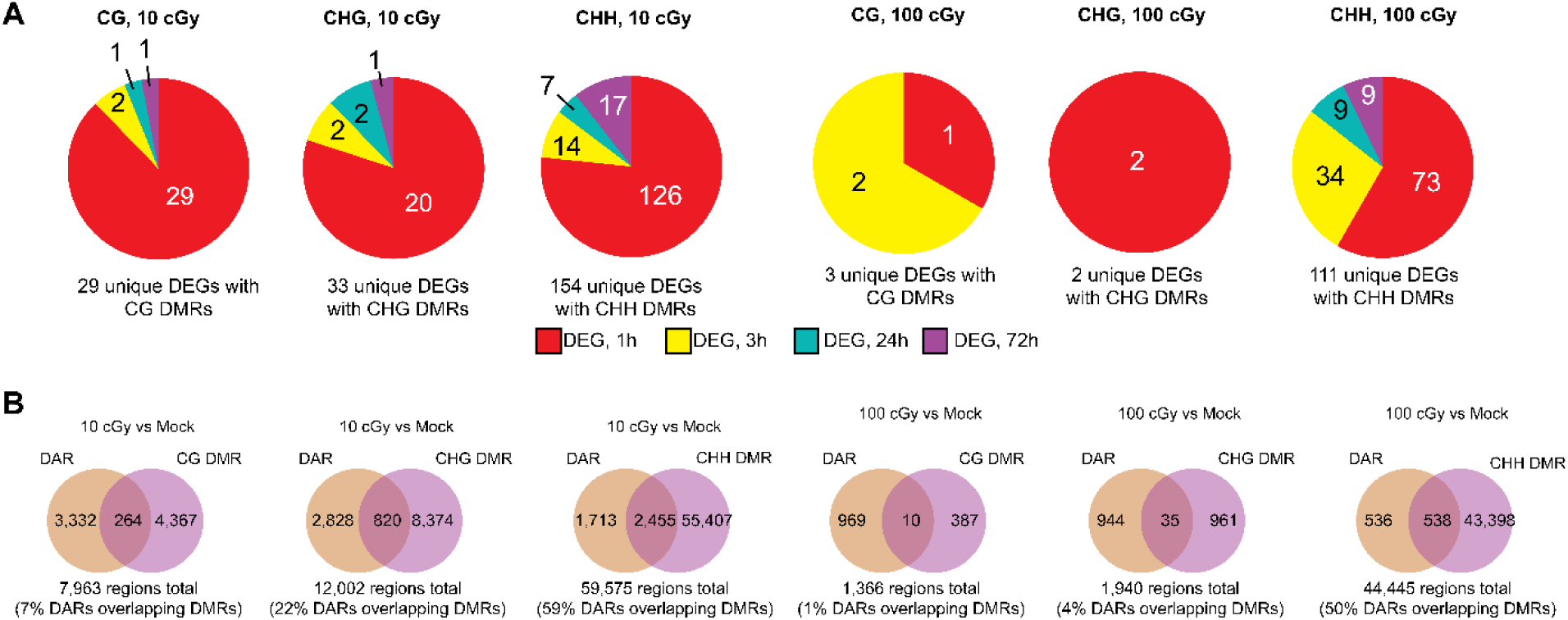
Comparison of methylation, accessibility, and gene expression changes in response to low dose ionizing radiation. (**A**) Distribution of differentially expressed genes (DEGs) with annotated differentially methylated regions (DMRs) by timepoint, ionizing radiation (IR) exposure, and site type. (**B**) Overlap between DMRs and differentially accessible regions (DARs) by IR exposure and site type.

To determine if the co-occurrence of differential methylation corresponded to changes in chromatin accessibility within genic regions, we plotted GC accessibility and endogenous methylation in genes (± 1 kb) with a proximal DAR (**Supplemental Fig. 4**). Decreased CG methylation but not CHG or CHH was observed at the TSS and corresponded with increased TSS accessibility at the TSS (**Supplemental Fig. 4**), and gene body methylation was largely CG, consistent with previous observations of methylation in *A. thaliana* (Cokus et al. 2008). Changes in CHH methylation in genes annotated with hyperaccessible regions appeared to be mostly small and confined to promoter regions (**Supplemental Fig. 4**). While there were fewer genes annotated with hypoaccessible regions, we found that endogenous methylation in these genes tended to be higher downstream of the gene body (**Supplemental Fig. 4**). We also plotted endogenous methylation in DARs annotated to genic regions and DARs annotated to intergenic regions (**Supplemental Fig. 5**). Endogenous methylation levels were generally higher in intergenic DARs compared to genic DARs regardless of methylation type (**Supplemental Fig. 5**). Changes in methylation in DARs were dependent on IR exposure and changes in GC accessibility; overall levels of GC accessibility across genic regions are consistently lower and closer to baseline in 100 cGy compared with 10 cGy. Compared to Mock-exposure, CHG and CHH methylation at 10 cGy was reduced in intergenic hyperDARs and unchanged in genic hyperDARs. There was also an increase in CHG and CHH methylation at 10 cGy within the central areas of genic hypoDARs but not in intergenic DARs. A smaller but similar increase in CHH methylation in genic hypoDARs was also seen for 100 cGy (**Supplemental Fig. 5**). This would suggest that changes in endogenous methylation do accompany changes in chromatin accessibility although the effect is specific to the level of IR exposure and the type of methylation and does not necessarily follow a similar pattern as changes in chromatin accessibility across the gene (**Supplemental Fig. 4 and 5**).

As an example of a gene that is differentially methylated and differentially accessible after exposure to IR, we plotted the endogenous methylation and GC accessibility for the gene *AT1G20860. AT1G20860* is differentially expressed 72 hr after 10 cGy IR exposure and has DMRs of each endogenous methylation type as well as a DAR (**Fig. 7**). *AT1G20860* consists of two exons and one intron. The changes in endogenous methylation are exclusively located within intron 1 close to the boundary with exon 2. A DAR was annotated to exon 2 and partly overlapped a ∼150 bp region of CHH differential methylation (**Fig. 7**). A closer examination of this region of CHH methylation revealed several DMRs of variable effect, although average CHH methylation across this region was ∼20% higher in plants exposed to 10 cGy than Mock-exposed plants (**Fig. 7**).

**Figure 7.**
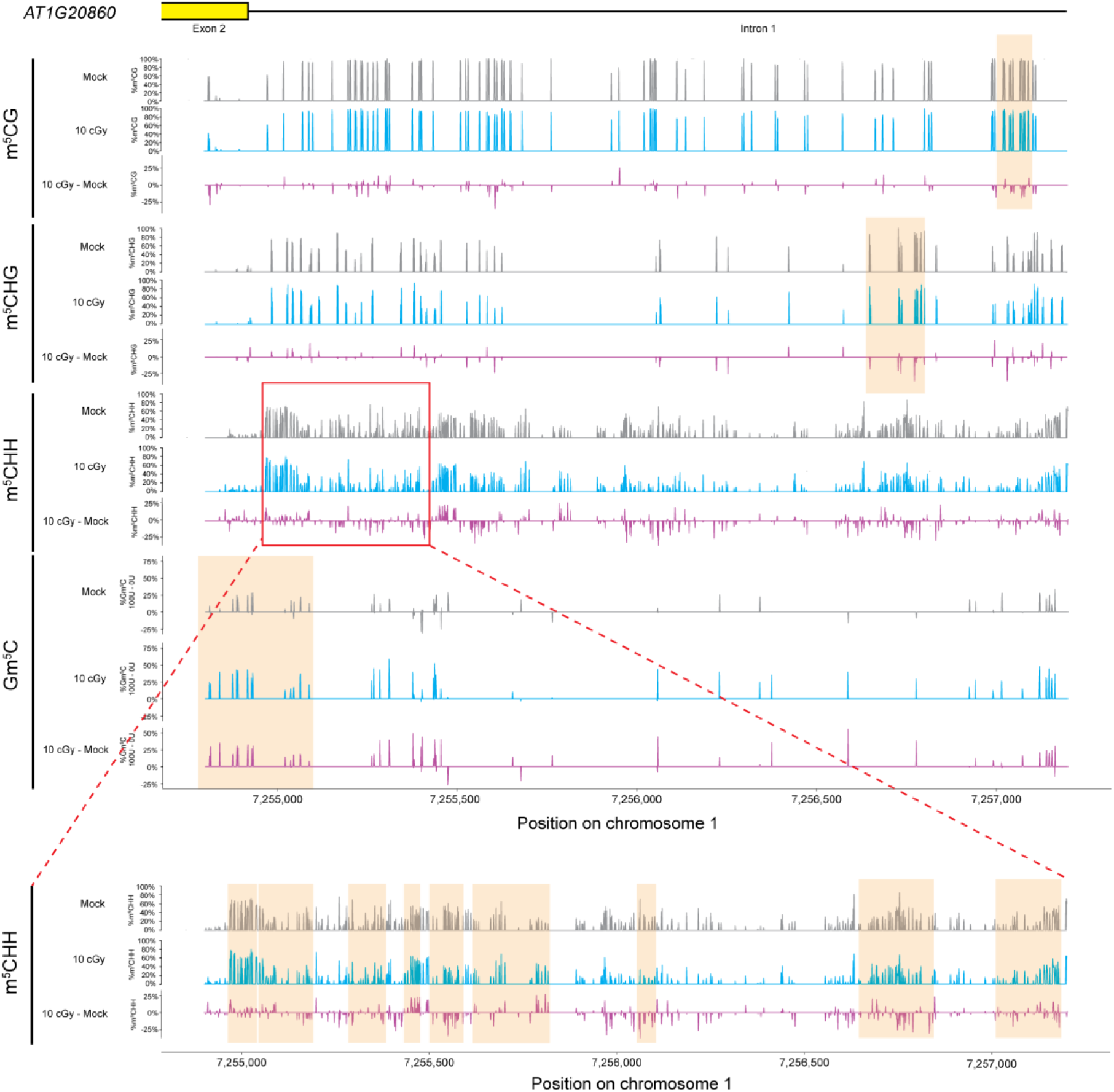
Differential methylation and differential accessibility in the *AT1G20860* gene. For each endogenous methylation type, the methylation for both 10 cGy (blue tracks) and Mock (grey tracks) exposures and the difference between them (purple tracks) are plotted. GC methylation is plotted as the difference between 100 U and 0 U M.CviPI for 10 cGy (blue tracks) and Mock (grey tracks) exposures and the difference between them (purple tracks). Regions of statistically significant difference are highlighted in orange. CHH DMRs for 10 cGy IR exposure are confined to a ∼500 bp region (red box) and zoom-in on this region to indicate CHH DMRs is provided.

As there are limited DE genes annotated with both DARs and DMRs, we also explored whether changes in endogenous methylation and chromatin accessibility were enriched near transposable elements (TE). As CHH methylation is known to be enriched near and may regulate the expression of TEs (e.g. (Martin et al. 2021), we sought to determine if epigenetic changes following IR exposure corresponded to changes in TE regulation. However, while most CHG and CHH DMRs were annotated to TE, there was only a limited association between methylation and chromatin accessibility in TE regions (**Supplemental Note 2**).

## Discussion

Plants first evolved when background levels of gamma radiation were higher than they are today (Karam and Leslie 1999). This likely shaped the biochemical pathways in plants that respond to abiotic stresses and protect the integrity of genomic DNA. Modern agriculture has employed ionizing radiation as a tool to generate novel varieties of plants with favorable traits and to sterilize fruits and vegetables (Deng et al. 2020; Lal et al. 2020; Ma et al. 2021). While these agricultural applications of IR to plants have typically employed relatively high doses, there is now increasing interest in the effects of lower doses of IR on plants, as might be encountered in the context of space travel or colonization of foreign worlds (De Pascale et al. 2021; Mohanta et al. 2021). In the current study, we have examined the joint effects of acute low-dose exposure to IR on gene expression, chromatin accessibility, and DNA methylation in the model plant, *A. thaliana*. Our goal was to gain insight into the nature and timing of genetic and epigenetic responses to IR in plants at these doses and to determine whether responses display specificity for IR as an abiotic stressor.

We observed robust transcriptional responses to acute doses of 10 and 100 cGy of IR, with significant changes in transcript levels detectable as soon as 1 hr after exposure. At least two temporal trajectories of response were detectable, an early response involving hundreds of DEGs and a later response involving fewer genes that were largely distinct from DEGs detected at earlier time points. DEGs, regardless of dose or time point, were much more frequently upregulated, rather than downregulated, in response to IR. Previous studies of the effects of high-dose IR on plants noted an enrichment for DEGs encoding products involved in sensing, signaling, or responding to DNA double-strand breaks (Culligan et al. 2006). At the lower doses employed here, where fewer DNA double-strand breaks per cell are induced, no such enrichment was observed. Rather, we observed significant evidence of activation of hormonal signaling pathways, in particular ethylene-activated signaling. Ethylene is well known to affect a broad array of abiotic stress responses in various plant species, including responses to salt, drought, heat, cold, light, and heavy metals (Anko et al. 2012). Consistent with this, there was also an enrichment for genes associated with iron homeostasis among genes DE 3 hr after IR exposure, which is known to be regulated by ethylene response factors (Yang et al. 2022). Interestingly, prior studies have noted that at high doses (hundreds to thousands of Gy), IR can inhibit fruit ripening due to effects on both the production of ethylene and response to its presence (Young 1965; Maxie et al. 1966). A prior microarray study of gene expression in *A. thaliana* exposed to an acute dose of 1 Gy noted that among significant DEGs were a set of hormone-responsive genes that included a number of ethylene-responsive genes (Kovalchuk et al. 2007).

Among the cellular effects of IR exposure is the production of ROS via the radiolysis of water. While toxic to cells, ROS can also play a significant signaling function. In plants, ROS are produced endogenously by enzymes resident in different tissues as well as by organelles such as chloroplasts as a byproduct of ATP production (Martin et al. 2022). There is clear evidence for cross-communication between the ethylene and ROS signaling pathways in multiple plant species responding to various abiotic stresses (Steffens 2014; Raza et al. 2022). In rice, ROS production in response to salt stress is dependent on ethylene signaling (Li et al. 2014). In *A. thaliana*, ethylene-insensitive mutants display high levels of oxidative stress in response to another abiotic stressor, drought (Cui et al. 2015). In addition, ethylene induces leaf chlorosis in responses to multiple stressors (Hodges and Coleman 1984; Niimi et al. 2015; Tao et al. 2015; Vaseva et al. 2021; Fatma et al. 2022). Thus, the activation of ethylene signaling pathways, as observed here for IR, is consistent with the chlorosis phenotype observed in this study (**Fig. 1B and C**) as well as findings for other abiotic stresses in plants and is likely mediated through the ability of IR to directly generate oxygen free radicals through its interaction with cellular water.

In order to assess the epigenetic response in *A. thaliana* to IR, we adapted and optimized a method, MAPit, previously developed and applied in mammalian cells, that allows for the simultaneous quantification of changes in chromatin accessibility and endogenous DNA methylation. Applying MAPit to *A. thaliana* plants, we observed a significant epigenetic effect of IR on chromatin accessibility, detecting more than 4,500 DARs that mapped to 1,267 genes. There was also a pronounced dose-dependent effect of IR on the directionality of changes in chromatin accessibility. At 10 cGy exposure, 99% of the DARs detected involved a shift towards greater accessibility. At 100 cGy, 79% of DARs displayed increased accessibility; most hypoDARs occurred in response to this higher dose.

The bias towards upregulation of DEGs in response to IR, particularly at the 10 cGy dose, together with the bias towards increased chromatin accessibility under the same conditions, suggests a model in which increased transcription factor binding drives the differences in gene expression observed after IR exposure. Consistent with this, a motif search of DARs identified a number of significantly enriched transcription factor binding sites, notably, multiple binding sites for transcription factors associated with ethylene signaling. These transcription factor binding sites were enriched among the promoter regions of DEGs responding to IR and the transcription factors, themselves, were also upregulated in response to IR. This concordance between the results obtained for gene expression and chromatin accessibility lends further support to the conclusion that ethylene-activated signaling plays a major role in the response to low-dose IR in *A. thaliana*.

DNA methylation is only one of several mechanisms that can act to regulate gene expression and, as such, might not be expected to display a tight correlation with the expression levels of individual genes. Indeed, observed changes in endogenous DNA methylation after IR exposure only broadly paralleled and did not show the same level of concordance as those observed for gene expression and chromatin accessibility. There was a robust response to IR exposure with more than 88,000 distinct regions of differential methylation detected (**Supplemental Fig. 2**), and with some dose dependence, as most DMRs were detected exclusively at the 10 cGy dose. The dose dependence of the endogenous methylation response was more pronounced when the sites of methylation were considered. Specifically, DMRs containing CG and CHG, while less frequent overall than CHH DMRs, were almost exclusively observed at the 10 cGy dose. For all sites, DMRs with CG, CHG, and CHH displaying reduced methylation predominated at the 10 cGy dose, the same dose at which most DEGs were upregulated. Integrative analysis of DARs and DMRs found that regions with decreased accessibility also corresponded to hypermethylation, especially of CHH motifs, regardless of the radiation application (**Supplemental Fig. 5**). We also observed enrichment, among DMRs, for binding motifs of several transcription factors associated with ethylene signaling. Of particular note is the ethylene response factor RRTF1, DNA recognition motifs for which were enriched in DAR and DMR sequences. RRTF1 is also known to regulate iron homeostasis in *A. thaliana* by repressing the expression of specific bHLH genes (Yang et al. 2022), two of which (*bHLH38*, *bHLH39*) are downregulated in response to IR at all time points assayed. Such results are broadly consistent with a model in which site-specific reductions in DNA methylation and derepression of gene expression correlate with increased chromatin accessibility.

From our data, we propose a possible mechanism of low-dose IR exposure in *A. thaliana* that is mediated by short-term induction and long-term regulation of the ethylene signaling response (**Fig. 8**). In this model, several genes involved in ethylene and other hormonal signaling responses (including some transcription factors) are quickly upregulated in response to IR, and associated signaling cascades are engaged. This is soon followed by altered regulation of iron homeostasis, potentially mediated by the ethylene response factors such as RRTF1 (as evidenced by the enrichment for their associated DNA-binding motifs in DMRs and DARs). Transcription largely returned to baseline at about 24 hr after the initial exposure. At 72 hr after the IR exposure, there is a second transcriptional response some of which includes ethylene response genes upregulated at 1 hr. We also see a robust epigenetic response at 72 hr, targeting sequences that are enriched for motifs associated with ethylene signaling and primarily consisting of increased chromatin accessibility and decreased methylation. We are not unable to ascertain when these changes occur. However, the largest and most consistent transcriptional response occurs early, and the concordance between IR doses with respect to DMRs and DARs, suggests that at least some of these epigenetic changes occur relatively soon after IR exposure, and they may accumulate and be sustained over time to regulate downstream responses.

**Figure 8.**
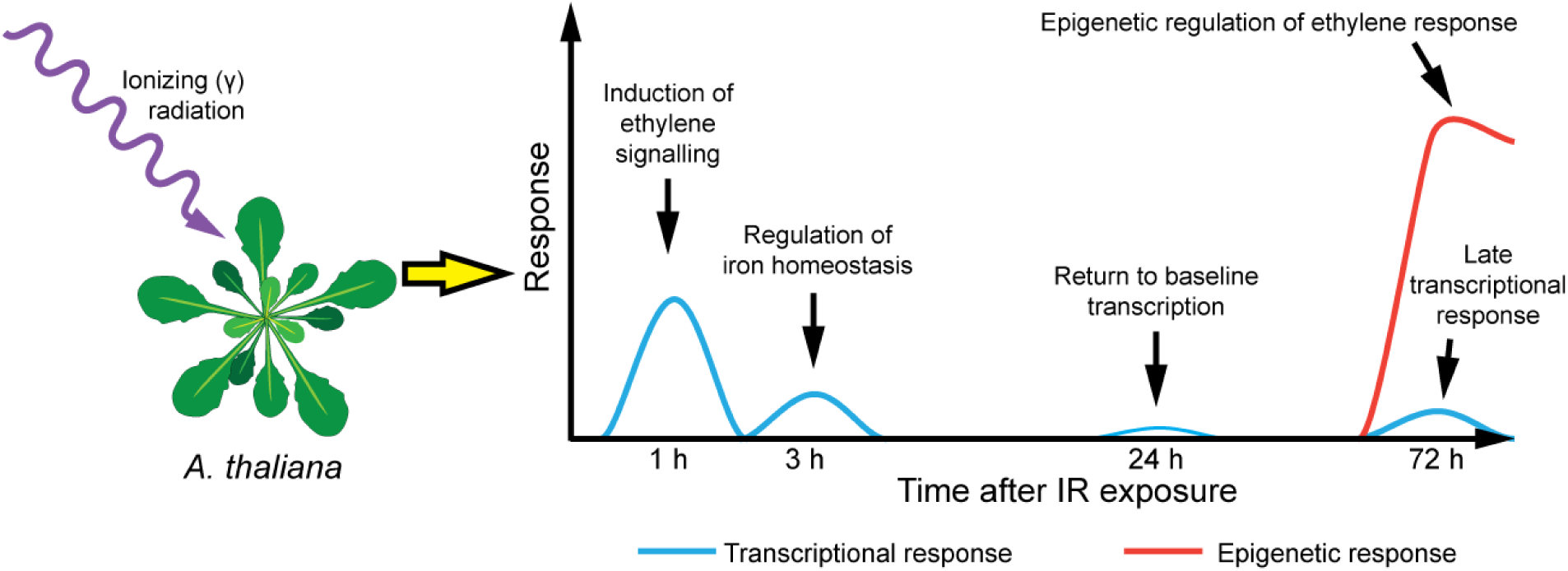
Proposed model of genetic response to low-dose IR in *A. thaliana*. Genes involved in ethylene and other hormonal signaling are induced within 1 hr after IR exposure. This is followed by expression of genes involved in regulating iron homeostasis by 3 hr, and largely a return to baseline transcription 24 hr after exposure. An additional, dose-dependent transcriptional response is observed at 72 hr after IR exposure, as is a robust epigenetic response, enriched for DNA sequences bound by ethylene response factors such as ERF1, RRTF1 and RAP2.3.

In summary, in response to acute exposures of as low as 10 cGy IR, *A. thaliana* mounts a transcriptional response that is rapid, detectable in as little as 1 hr, and robust, with detection of >650 significant DE genes strongly biased towards induced gene expression. These significantly upregulated DE genes are associated with increases in chromatin accessibility. The common feature of both of these responses is activation of the ethylene signaling pathway, suggesting that this is the primary driver of this stress response. Significant changes to DNA methylation are also observed in response to low-dose IR. Some of the hypomethylated sites detected in response to IR occur within binding motifs for ethylene-activated transcription factors, consistent with the findings from transcriptional and chromatin accessibility measures. Our multi-omics approach, simultaneously measuring and then jointly analysing gene expression, chromatin accessibility and endogenous DNA methylation, facilitated by our adaptation of the MAPit technique to plants, was key to uncovering this relationship between ethylene signaling and the low-dose IR response in *A. thaliana*. Overall, our findings are consistent with a model in which *A. thaliana* responds to low-dose IR by activating a commonly used stress response pathway through a combination of derepression via reductions in site-specific DNA methylation and increases in both chromatin accessibility and gene transcription at genes related to ethylene signaling.

## Methods

### Experimental design

*Arabidopsis thaliana* (Col-0 ecotype) seeds were sown in autoclaved soil and cold-treated at 4°C for 3 days to promote uniform germination (Zhou et al. 2017). Plants were grown under a short-day condition (8 hr light, 16 hr dark) at 22°C for 1 week, followed by transplantation of each seedling to individual cells in 36-cell trays. At 4 weeks of growth, plants were placed in a ^137^Cs irradiator and exposed to gamma IR at 1.4 cGy per second. Plants were irradiated for 7 sec (10 cGy equivalent), 71 sec (100 cGy equivalent) or not exposed (0 cGy or “Mock”). Whole rosettes were collected at 1 hr, 3 hr, 24 hr, and 72 hr post-exposure for RNA-seq and at 72 hr post-exposure for MAPit (**Fig. 1A**). RNA-seq was performed with three biological replicates per condition, whereas epigenetic studies were conducted with two biological replicates.

### RNA sequencing

Total RNA was extracted from approximately 0.1 g whole rosette tissues using the RNeasy Plant Mini Kit (Qiagen). RNA samples were quantified using a NanoDrop spectrophotometer (NanoDrop Technologies) and quality was assessed using an Agilent 2100 Bioanalyzer. RNA-seq library construction was performed by the University of Florida Interdisciplinary Center for Biotechnology Research (UF-ICBR, RRID:SCR_019152), using poly(A) selection and the NEBNext Ultra II Directional RNA Library Prep Kit (New England Biolabs). RNA-seq (2 × 100 nt paired-end reads) was performed using a HiSeq 3000 instrument (Illumina), yielding an average of 38 million reads/sample.

### RNA-seq alignments and quantification

Raw RNA-seq reads were trimmed using Trimmomatic (version 0.32; (Bolger et al. 2014) to remove adapters and low-quality reads. A Phred-33 quality score of 20 was used for quality trimming of ends of reads (parameters *LEADING:20 TRAILING:20*). Sliding-window trimming was also performed with a window size of 4 nt and a minimum average quality of 15 (parameter *SLIDINGWINDOW:4:15*). Reads that were less than 80 nt after trimming were removed from further analysis (parameter *MINLEN:80*).

Trimmed reads were then aligned paired-end to the *A. thaliana* TAIR10 genome assembly (http://www.arabidopsis.org/; (Lamesch et al. 2012) using the STAR aligner (version 2.5.2b; (Dobin et al. 2013), with the following parameters: *--runThreadN 8 --outSAMtype BAM SortedByCoordinate --limitBAMsortRAM 48000000000 --outFilterType BySJout --outFilterMultimapNmax 20 --alignSJoverhangMin 8 --alignSJDBoverhangMin 3 --alignIntronMin 7 --alignIntronMax 40000 --alignMatesGapMax 40000 --outSAMattributes Standard --outFilterIntronMotifs RemoveNoncanonical --outSAMstrandField intronMotif --quantMode TranscriptomeSAM*. Resulting alignments were then passed to RSEM (version 1.2.28; (Li and Dewey 2011) using the *rsem-calculate-expression* program with parameter *--alignments* to estimate the abundances of TAIR10-annotated transcripts and genes.

### Gene expression analysis

Gene expression abundance estimates were expressed as copies per million (CPM), calculated as:

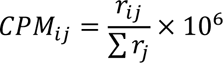

where *r_ij_* is the number of reads mapped to gene *i* in sample *j* and Σ*r_j_* is the sum of all mapped reads for sample *j*. A gene was considered detected if it had CPM > 0 in at least 2 biological replicates for a given treatment group. The data were then upper-quartile normalized and log_2_ transformed (Bullard et al. 2010; Dillies et al. 2013).

The mixed effects model, *Y_ijk_ = μ + d_i_ + t_j_ + d_i_t_j_ + ε_ijk_*, was fit separately for each gene, where *Y_ijk_* is the log_2_-transformed normalized CPM, *d_i_* is IR dosage (*i* = Mock, 10 cGy, or 100 cGy), *t_j_* is the time point (*j* = 1 hr, 3 hr, 24 hr, or 72 hr), and *k* is the individual sample. All variables were treated as fixed effects. Residuals (ε) were assumed to be distributed N(0, σ*_n_*). Individual comparisons of irradiated samples to mock-treated samples were calculated for each time point (10 cGy vs Mock for 1 hr, 3 hr, 24 hr, or 72 hr; 100 cGy vs Mock for 1 hr, 3 hr, 24 hr, or 72 hr). A false discovery rate (FDR) correction was applied (Benjamini and Hochberg 1995; Verhoeven et al. 2005) and FDR-corrected *P* < 0.05 were considered to be significant. Significant genes with a log_2_ fold change (log_2_ FC) ≥ 1 and log_2_ FC ≤ −1 were considered to be up- and downregulated (relative to mock treatment), respectively.

Changes in log_2_ FC over time of differentially expressed genes (DEGs) were modelled using the mixed effects model *Y_ijk_ = μ + c_i_ + t_j_ + c_i_t_j_ + ε_ijk_*, where *Y_ijk_* is log_2_ FC, *c_i_* is comparison (*i* = 10 cGy vs Mock, 100 cGy vs Mock), *t_j_* is time point (*j*=1 hr, 3 hr, 24 hr, 72 hr), *c_i_t_j_* is the interaction between comparison and timepoint (the predictor term of interest), and *k* is the individual sample. All variables were treated as fixed effects. Residuals (ε) were assumed to be distributed N(0, σ*_n_*). Gene ontology enrichment analysis was performed on the sets of DEGs to infer which biological processes were perturbed following irradiation (Ashburner et al. 2000; The Gene Ontology Consortium 2019). Gene ontology annotations were downloaded from www.arabidopsis.org. For each treatment comparison (10 cGy vs Mock; 100 cGy vs Mock) and each time point, gene ontology term enrichment was performed by comparing the set of up- or downregulated genes (absolute log_2_ FC ≥ 1 and FDR-corrected *P* < 0.05) to the set of non-up- or non-downregulated genes (absolute log_2_ FC ≤ 1 or FDR-corrected *P* ≥ 0.05). *P* values were calculated using a Fisher’s exact test (FET) as implemented in the “Gene Set Enrichment” procedure in JMP Genomics 9 (SAS Institute).

### Methyltransferase accessibility protocol for individual templates (MAPit)

Details of how we adapted the MAPit protocol for application to plants, including optimization of the protocol for application to plant tissue can be found in **Supplemental Note 1**. Approximately 0.5 g of whole rosette tissues from two different groups of ∼72 *A. thaliana* seedlings 72 hr post-exposure (Mock, 10 cGy, or 100 cGy) were ground separately in a pre-chilled mortar with a pestle on ice in 5 ml of nuclei isolation buffer (15 mM Tris, pH 7.8, 0.30 M sucrose, 0.3 mM spermine, 0.125 mM spermidine, 20 mM NaCl, 80 mM KCl, 15 mM β-mercaptoethanol, 0.1 mM EGTA, 1 mM EDTA, 1/100 volume of protease inhibitor cocktail (C0001, TargetMol, Boston, USA), 0.15% Triton X-100). Ground tissue was then sequentially filtered with four layers of cheesecloth, one layer of Miracloth, and one 105-µm mesh polypropylene screen. Nuclei were pelleted by centrifugation at 1500*g* for 10 min at 4°C and washed using cell resuspension buffer (CRB) (20 mM HEPES, pH 7.5, 70 mM NaCl, 0.25 mM EDTA, 0.5 mM EGTA, 0.5% glycerol, 10 mM DTT, 0.25 mM PMSF). Two reactions containing ∼ 3 million nuclei (prepared from separate sets of rosettes) each were incubated in parallel with 100 U or 0 U of M.CviPI (Xu et al. 1998) (New England Biolabs) in methylation buffer (320 µM *S*-adenosyl-*L*-methionine in CRB) for 15 min at 37°C. The 0 U M.CviPI reaction is required for MAPit of *Arabidopsis* chromatin to correct for endogenous CG, CHG, and CHH methylation, where H is A, C, or T. The reaction was terminated by addition of an equal volume of methylation stop buffer (100 mM NaCl, 10 mM EDTA, 1% SDS). Nucleic acids were treated with 10 µg/ml RNase A for 30 min at 37°C followed by incubation with 100 µg/ml proteinase K overnight at 50 °C. Genomic DNA was extracted using phenol-chloroform-isoamyl alcohol (25:24:1, v/v) phase separation, followed by ethanol precipitation and resuspension in molecular biology-grade, sterile water.

### Whole genome bisulfite sequencing

Whole genome bisulfite sequencing was performed as previously described (Zhou et al. 2019). Approximately 400 ng of genomic DNA from *A. thaliana* 72 hr post-exposure (Mock, 10 cGy, or 100 cGy) was processed for bisulfite sequencing library construction using NEBNext Ultra II DNA Library Prep Kit (NEB) along with methylated and indexed NEXTflex Bisulfite-seq Barcodes (Bio scientific). Bisulfite conversion was carried out using the EZ DNA Methylation-Gold Kit (Zymo Research). Sequencing was performed on a HiSeq 3000 platform using 2 × 101 cycle multiplex paired-end reads. The raw sequencing reads were trimmed using Trimmomatic (version 0.39) and subjected to quality control using FastQC (version 0.11.7), then aligned to the *Arabidopsis* TAIR10 reference using BSMAP (version 2.87) (Xi and Li 2009). Mapping statistics for whole genomic bisulfite sequencing are presented in **Supplemental Table 6**. Methylation calling was performed with CSCALL (version 1.0; http://compbio.ufl.edu/cscall), which is a part of DMAP2 pipeline (Ianov et al. 2017; Zhou et al. 2019). The non-bisulfite conversion rate together with the sequencing error rate of thymidine to cytosine at cytosine positions was estimated by calculating the average methylation at HCHH sites in the chloroplast genome in M.CviPI-treated samples (estimated bisulfite conversion rate = 96.4%). A cytosine site was included in subsequent analysis if it reached at least 10x read coverage across both replicates in a given treatment.

### Differential chromatin accessibility analysis

M.CviPI-induced methylation at GC sites was used to assess changes in chromatin accessibility following exposure to IR. GC sites were retained if they had at least 10 reads of coverage summed across both replicates. Due to efficiency differences in GC methylation rates across samples, we employed the following normalization approach for the 100U samples. Replicates were first globally normalized for differences in the total numbers of reads per sample. A scale factor was calculated as the total number of reads in each replicate divided by the average total reads across replicates. The scale factor was then applied to both the total counts and the methylated counts. Sites were then split into five equally sized groups based on their average methylation rate and a group-specific scale factor was calculated to normalize the methylated counts. Within each group, a scale factor for each replicate was computed as the total methylated reads across all sites divided by the average total methylated reads across replicates.

Chromatin accessibility 72 hr post-exposure was determined by first correcting for background endogenous methylation of CG, CHG, and CHH in *A. thaliana*. To do so, the difference in GC methylation was calculated between the combined duplicate samples treated with 100 U and 0 U of M.CviPI in each condition (i.e., 10 cGy 100 U - 0 U, 100 cGy 100 U - 0 U, and 0 cGy 100 U - 0 U) (Zhou et al., in preparation). A quantitative estimate of GC accessibility was also calculated as the difference in GC methylation between 10 cGy or 100 cGy and Mock, corrected for endogenous methylation ((i.e., 10 cGy or 100 cGy (100 U M.CviPI treated - 0 U) - Mock (100 U M.CviPI treated - 0 U)). Fisher’s exact test (FET) was employed to identify individual sites of accessible chromatin by testing differential GC methylation between 100 U and 0 U of M.CviPI for each condition (Mock vs 10 cGy or Mock vs 100 cGy), FDR-adjusting *P* values. GC sites indicative of loci with significant changes (FET FDR *P* < 0.05) in GC methylation after the addition of 100 U M were considered consistent with loci of open (increased GC methylation) vs closed (decreased GC methylation) chromatin and were analyzed further. Differential accessibility of individual GC sites was tested using a Cochran-Mantel-Haenszel (CMH) test stratified by units of M.CviPI. A GC site was considered differentially accessible if it had an FDR-corrected CMH *P* < 0.05 for Mock-corrected GC methylation at 10 cGy or 100 cGy and ≥ 10% difference in methylation level between conditions. Individual GC sites indicative of active chromatin were grouped into the same region if they were spaced ≤ 100 bp apart and the difference in GC accessibility was in the same direction (e.g., both displaying increased or decreased accessibility; **Supplemental Fig. 6,** for example). Regions were considered further if there were at least 5 GC sites consistent with loci of open or closed chromatin.

Differentially accessible regions (DARs) were defined as regions containing at least two GC sites differing from the Mock reference, where the differences were in the same direction. Several approaches to quantifying differential accessibility were tested (**Supplemental Note 3**). For each region, a binomial logistic regression with the terms IR exposure (Mock vs 10 cGy or Mock vs 100 cGy), M.CviPI units (0 U, 100 U), and the interaction term IR exposure × M.CviPI units. Coverage at individual sites within a region was accounted for by fitting a random intercept for each site within a region. The IR exposure × M.CviPI units interaction was used as the predictor variable for differential accessibility and *P* values were FDR adjusted. A region was considered a DAR if it met one of two criteria: (1) a region must have at least 2 GC sites with an FDR-corrected CMH *P* < 0.05 and ≥10% difference in GC accessibility; or (2) a region has a binomial regression FDR-corrected *P* < 0.05 and an average GC accessibility difference between IR exposed and Mock ≥10%.

### Differential endogenous methylation analysis

Differential methylation of individual CG, CHG, and CHH sites between irradiated (10 cGy or 100 cGy) and mock-irradiated (Mock) were determined for nuclei treated with 0 U M.CviPI, i.e., using a FET (Sun et al. 2014). Significance *P* values were adjusted for multiple comparison using the FDR method (Benjamini and Hochberg 1995). A cytosine was considered differentially methylated between conditions if it had an FDR-corrected FET *P* < 0.05 and ≥ 10% difference in methylation level between conditions. For each assayed endogenous methylation type (CG, CHG, and CHH), the same grouping strategy used for GC sites was employed: sites were grouped into regions if they were ≤100 bp apart and the difference in methylation between exposed and Mock-exposed was in the same direction (**Supplemental Fig. 6**). Like differential accessibility, several models for defining differential methylation were tested (**Supplemental Note 3**). Ultimately, for each region, a binomial logistic regression with IR exposure as the predictor term (Mock, 10 cGy or 100 cGy) was fit. Coverage at individual sites within a region was accounted for by fitting a random intercept for each site within a region. Significance *P* values were FDR adjusted and an FDR-corrected *P* < 0.05 was considered statistically significant. Endogenous, differentially methylated regions (DMRs) were identified by one of two criteria: (1) regions containing at least two methylated sites (CG, CHG, or CHH) with FDR-corrected FET *P* < 0.05 and a ≥10% difference in average methylation levels between exposed and mock-treated samples across the region; or (2) an FDR-corrected binomial regression *P* < 0.05 and a ≥10% difference in average methylation levels between exposed and mock-treated samples across the region.

### DNA-binding site motif analysis

Detection of known protein binding motifs within DARs and DMRs was performed using the MEME suite of DNA motif analysis tools (Bailey et al. 2015). The BEDtools (Quinlan and Hall 2010) function ‘getfasta’ was used to extract DNA sequences using a BED file containing the TAIR10 genomic coordinates of DARs or DMRs as input. Queried regions <12 bp were centered and extended to 6 bp on both sides (i.e., 12 bp total) to search for potential motifs that overlapped with the regions. Repetitive sequences and low complexity sequences in the set of extracted DAR and DMR sequences were then masked by RepeatMasker (http://www.repeatmasker.org) (Smit et al. 2013). The masked sequences were used as input for Simple Enrichment Analysis (Bailey and Grant 2021) and compared to two databases of *Arabidopsis* DNA recognition motifs: DAP-seq targets from O’Malley et al. (O’Malley et al. 2016) and motifs from protein-binding microarrays from Franco-Zorrilla et al. (Franco-Zorrilla et al. 2014).

### Code availability

All code pertaining to the analysis of the data presented in this study is available on GitHub at https://github.com/jrbnewman/Arabidopsis_radiation.

## Data Access

The RNA and whole genome bisulfite sequencing data generated in this study have been submitted to the NCBI BioProject database (https://www.ncbi.nlm.nih.gov/bioproject/) under accession number PRJNA681215.

## Competing interests statement

The authors declare that they have no competing interests.

## Acknowledgments

This work was supported by funding from the Defense Threat Reduction Agency (HDTRA1-16-0048).

## Notes

### Competing Interest Statement

The authors have declared no competing interest.

### Summary of Updates

Reformatted manuscript for Genome Research submission

