## Supplemental Information for "Multi-omics profiling reveals ethylene signalling as a key pathway underlying both genetic and epigenetic responses to low-dose ionizing radiation in *Arabidopsis*"

**Main text Supplemental tables and figures**

**Supplemental Table 1.** Gene Ontology enrichment of differentially expressed genes

File: Supplemental_Table_1_RNAseq_arabidopsis_GO_results.xlsx

**Supplemental Table 2.** Transcription factor DNA recognition motif analysis of differentially accessible regions

File: Supplemental_Table_2_DAR_motif_enrichment.xlsx

**Supplemental Table 3.** Transcription factor DNA recognition motif analysis of promoter regions of differentially expressed genes

File: Supplemental_Table_3_Motif_analysis_DEG_promoters.xlsx

**Supplemental Table 4.** Expression of genes associated with DNA-binding motifs overrepresented in differentially accessible regions

File: Supplemental_Table_4_Expression_of_genes_assoc_DAR_motifs.xlsx

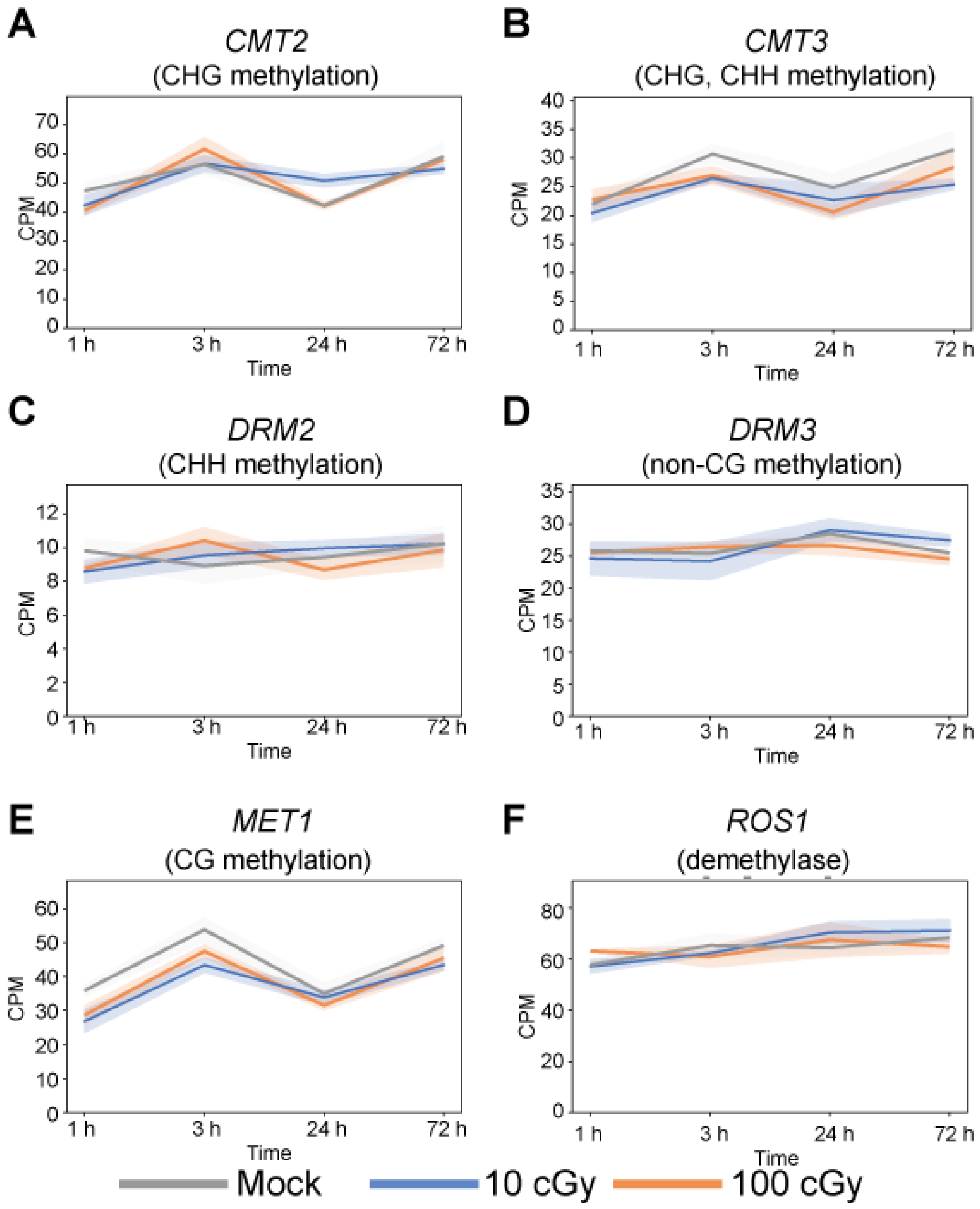

**Supplemental Figure 1.** Expression of transcribed DNA methyltransferases at 1 h, 3 h, 24 h, and 72 h after exposure to ionizing radiation exposure (**A**) *CMT2*, (**B**) *CMT3*, (**C**) *DRM2*, (**D**) *DRM3*, and (**E**) *MET1*. (**F**) Expression of the *ROS1* demethylase gene at 1 h, 3 h, 24 h, and 72 h after ionizing radiation exposure.

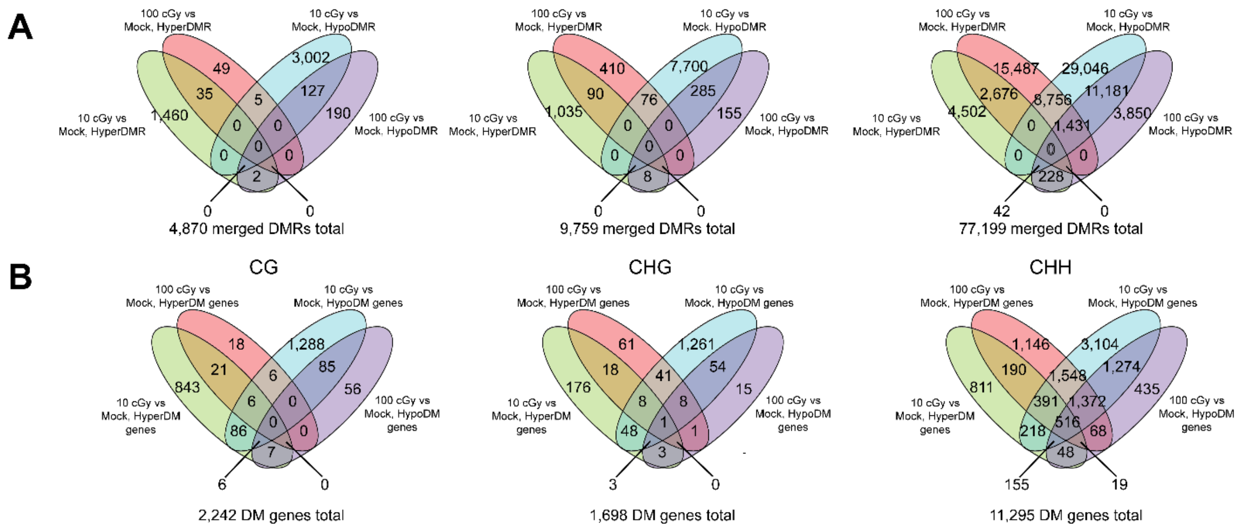

**Supplemental Figure 2.** Overlap of differentially methylated regions and genes. (A) Overlap of differentially methylated regions (DMRs) segregated as hypermethylated regions (hyperDMRs) and hypomethylated regions (hypoDMRs) regions (DMRs) by site type. (B) Overlap of differentially methylated (DM) genes segregated as hypermethylated (hyperDM) and hypomethylated (hypoDM) genes by site type.

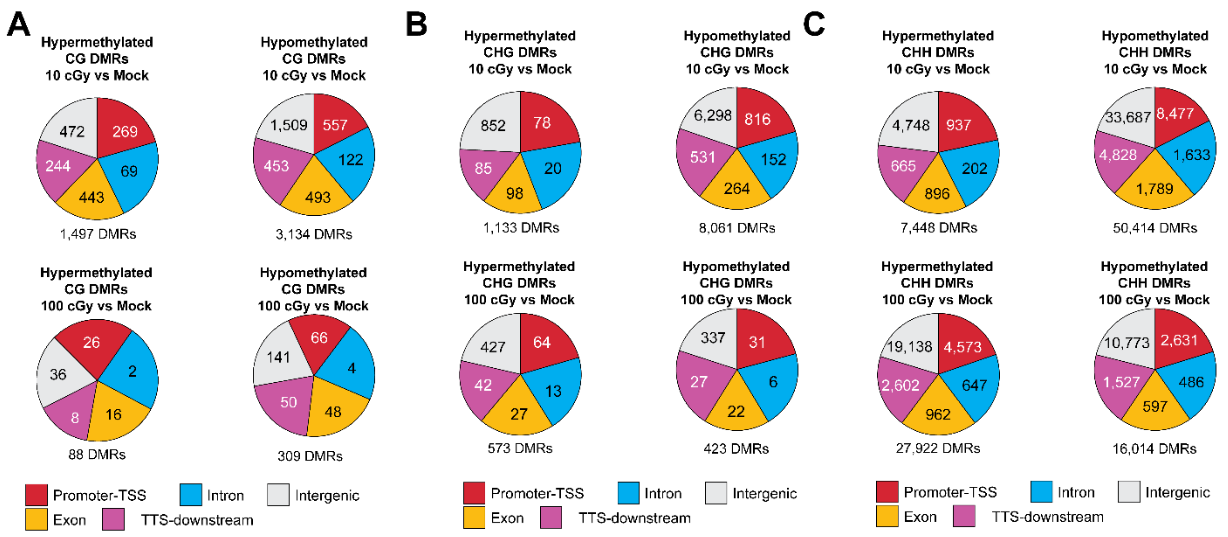

**Supplemental Figure 3.** Distribution of genic features annotated to differentially methylated regions. (**A**) CG differentially methylated regions (DMRs) (**B**) CHG DMRs (**C**) CHH DMRs. Proportion of pie chart is normalized to the total number of CG, CHG, or CHH motifs within all DMRs of that feature type.

**Supplemental Table 5.** Transcription factor DNA recognition motif analysis of differentially methylated regions

File: Supplemental_Table_5_DMR_motif_enrichment.xlsx

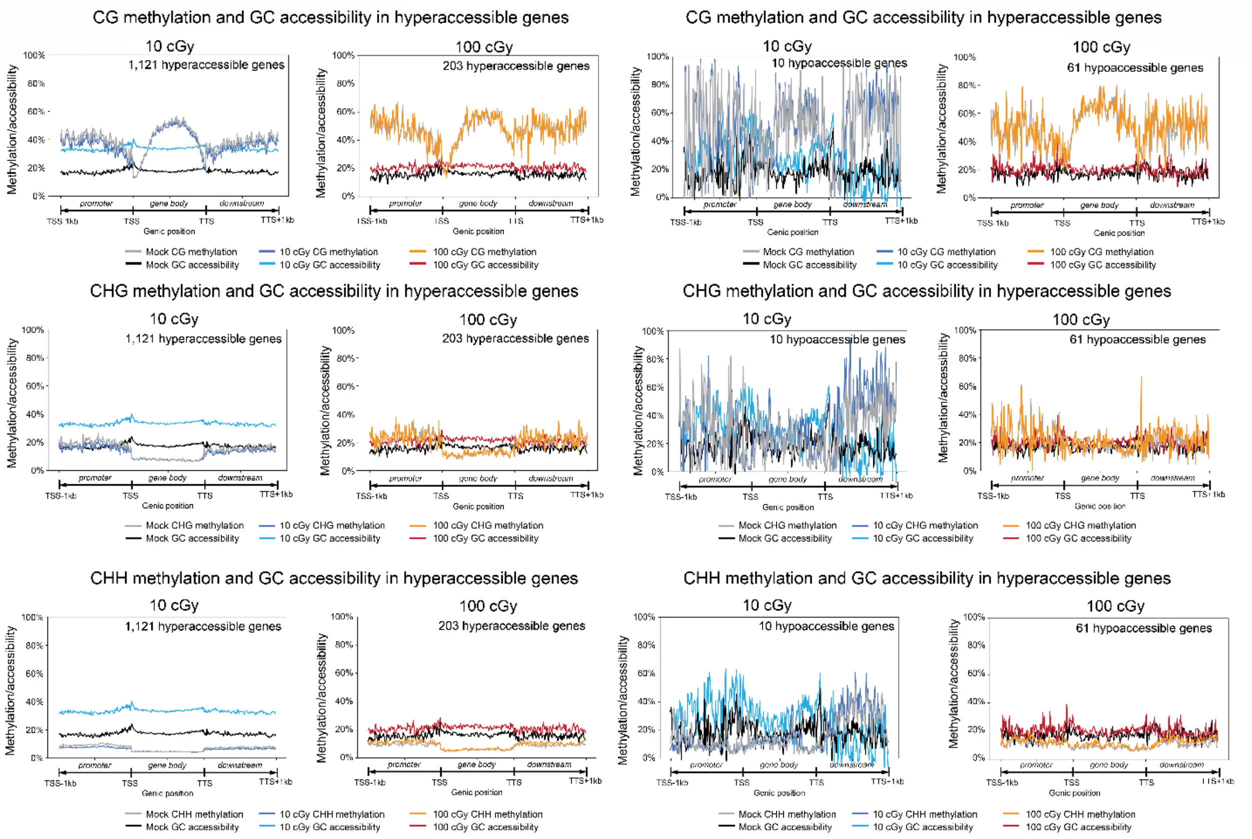

**Supplemental Figure 4.** Endogenous methylation and chromatin accessibility in gene bodies. Endogenous methylation (CG, CHG, CHH) and GC accessibility in genes annotated to hyperaccessible and hypoaccessible DARs by ionizing radiation exposure (10 cGy, 100 cGy). TSS = transcriptional start site; TTS = transcriptional termination site.

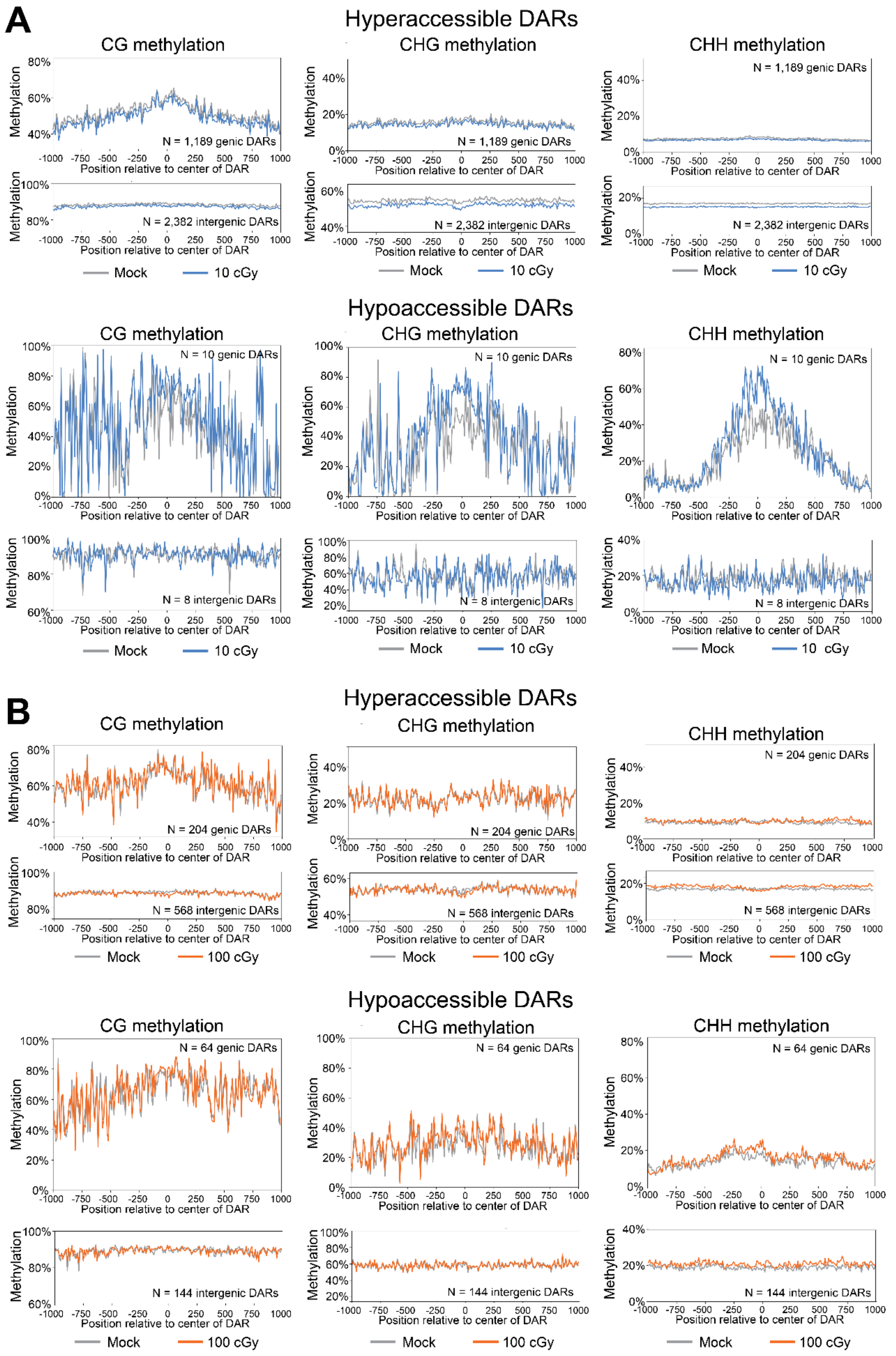
A
Hyperaccessible DARs
CG methylation
CHG methylation
CHH methylation
Methylation
Position relative to center of DAR
N = 1,189 genic DARs
N = 2,382 intergenic DARs
Mock
10 cGy
Hypoaccessible DARs
N = 10 genic DARs
N = 8 intergenic DARs
B
Hyperaccessible DARs
N = 204 genic DARs
N = 568 intergenic DARs
Mock
100 cGy
Hypoaccessible DARs
N = 64 genic DARs
N = 144 intergenic DARs

**Supplemental Figure 5.** Endogenous methylation in differentially accessible regions. Endogenous methylation (CG, CHG, CHH) in hyperaccessible and hypoaccessible differentially accessible regions (DARs) for (**A**) 10 cGy and (**B**) 100 cGy.

**Supplemental Table 6.** Whole genome bisulfite sequencing mapping statistics

File: Supplemental_Table_6_WGBS-MAPit_mapping_statistics.xlsx

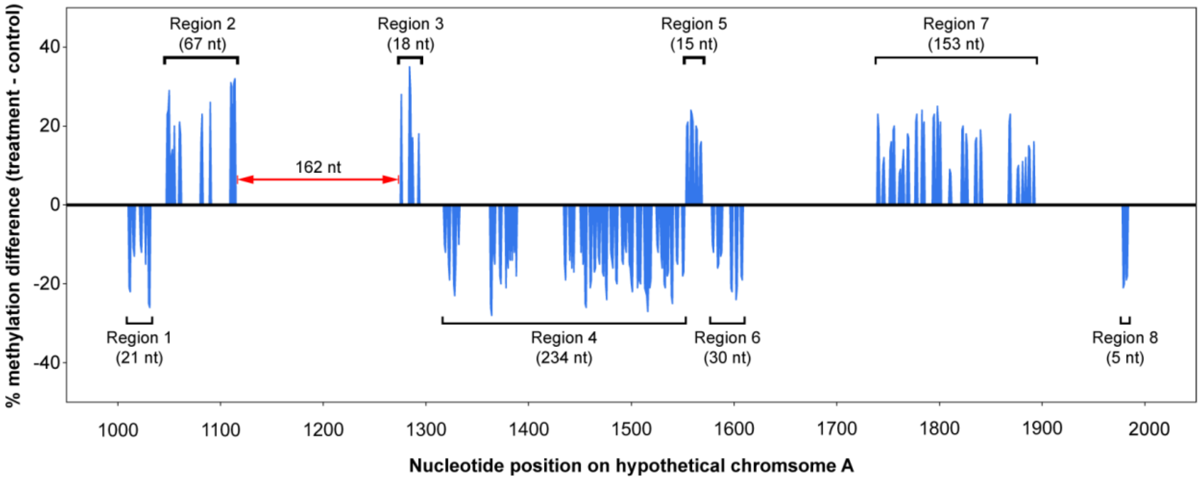

**Supplemental Figure 6.** Strategy employed in grouping methylation sites into regions. Methylation sites on hypothetical chromosome A are incorporated into the same region if (1) the methylation site was the same (CG, CHG, CHH, GC), (2) the difference in methylation level between exposed and mock conditions were in the same direction (i.e. hypermethylation, hypomethylation) and were spaced no more than 100 nt apart. Methylation sites less than 100 nt apart and had a difference in methylation level in opposite directions were considered to be in separate regions, as were sites that were more than 100 nt and had a difference in methylation level in same directions (example indicated in red).

**Supplemental Note 1.** Optimization of of M.CviPI titer for application of methyltransferase accessibility protocol for individual templates (MAPit) in plants

### Methods

Data used for the optimization of the MAPit protocol was provided by Zhao et al (unpublished results).

### Plant materials and treatments

*Arabidopsis thaliana* (Columbia ecotype, Col) seeds were sterilized with 10% bleach and 70% ethanol, then sown on the surface of solid 0.5x MS nutrient medium (Zhou et al. 2017). Plants were grown in a long-day condition (16 h light/8 h dark) at 22°C for 7 days. On day 8, 4 hr after the photoperiod began, whole seedlings were harvested and immediately prepared for MAPit of nuclei obtained from fluorescence-activated nuclei sorting (FANS) or tissue grinding. Treatments were done as two independent biological replicates.

### Fluorescence-activated nuclei sorting (FANS) for plant samples

Approximately 0.3 g of whole seedlings were collected and immediately minced in 3 ml of pre-chilled nuclei isolation buffer for FANS (15 mM Tris, pH 7.8, 0.3 mM spermine, 0.125 mM spermidine, 20 mM NaCl, 80 mM KCl, 15 mM β-mercaptoethanol, 0.1 mM EGTA, 1 mM EDTA, 0.15% (v/v) Triton X-100). The slurry was filtered through a four-layer bundle of cheesecloth and then one layer of Miracloth. The nuclei were then centrifuged at 2,200 *g* for 20 min at 4°C and the pellets were resuspended in 500 µl isolation buffer pre-chilled to 4°C. Nuclei sorting was executed as previously described medium (Vrána et al. 2012). In brief, crude nuclei were stained with 2 μg/ml of 4,6-diamidino-2-phenylindole (DAPI) in the dark for 20 min and sorted on a BD FACSAria-II SORP (BD Biosciences) flow cytometer. Blue laser (488 nm, 100 mW) was used for forward *versus* side scatter (FSC *vs*. SSC) gating, and UV laser (355 nm, 100 mW) was used for the excitation of DAPI fluorescence. Data acquisition and analysis were performed with BD FACSDiva software (BD Biosciences).

**Methyltransferase accessibility protocol for individual templates (MAPit) for plants**

MAPit was performed on both fresh tissues and FANS samples. For MAPit of fresh tissues, approximately 0.5 g 7-day-old *Arabidopsis* seedlings were ground in a pre-chilled mortar with pestle on ice in 5 ml of Nuclei Isolation Buffer (NIB; 15 mM Tris, pH 7.8, 0.30 M sucrose, 0.3 mM spermine, 0.125 mM spermidine, 20 mM NaCl, 80 mM KCl, 15 mM β-mercaptoethanol, 0.1 mM EGTA, 1 mM EDTA, 1/100th volume of protease inhibitor cocktail (catalog no. C0001; Targetmol, Boston, USA), 0.15% (v/v) Triton X-100). Next, the slurry was filtered sequentially once through a four-layer bundle of cheesecloth, one layer of Miracloth, and then one 105-μm mesh polypropylene screen. Nuclei were pelleted by centrifugation at 1,500 *g* for 10 min at 4°C. For FANS samples, the sorted nuclei were centrifuged at 1,000 *g* for 10 min at 4°C. The pellets from fresh tissues or FANS samples were washed in cell resuspension buffer (CRB; 20 mM HEPES, pH 7.5, 70 mM NaCl, 0.25 mM EDTA, 0.5 mM EGTA, 0.5% glycerol (v/v), 10 mM DTT (freshly added), and 0.25 mM PMSF (freshly added). Nuclei were then probed for accessible GC sites in methylation buffer (320 μM *S*-adenosyl-*L*-methionine in CRB) with 100 units (U) of M.CviPI (New England Biolabs, Ipswich, USA) for 15 min at 37°C. Methylation reactions were terminated by addition of an equal volume of methylation stop buffer (100 mM NaCl, 10 mM EDTA, 1% SDS). Nuclei were treated with RNase A for 30 min at 37°C and deproteinated by incubation with 100 µg/mL proteinase K overnight at 50°C. Genomic DNA was extracted using phenol-chloroform-isoamyl alcohol (25:24:1 (v/v)) phase separation, followed by ethanol precipitation, and resuspension in dd$H_2O$. Detailed steps of the procedure and supporting notes are provided in the Supplemental Protocol below.

### RESULTS

#### Overview of MAPit workflow in plant materials

In MAPit, C-5 DNA methyltransferases (DMNTs) are used as probing agents to enter nuclei and modify accessible DNA sites in open regions of chromatin (**Supplemental Fig. 7**) (Pondugula and Kladde 2008; Pardo et al. 2011; Pardo et al. 2015). Subsequent bisulfite sequencing provides a single-nucleotide-resolution readout of the methylation status of all C residues as either unmethylated or methylated ($m^5C$) by endogenous DNMTs or the exogenously supplied DNMT. The mapped patterns of exogenous $m^5C$ and nuclease cutting provide analogous outputs, namely, averaged chromatin accessibility used to infer the positions of nucleosomes and DNA-bound transcription factors along chromatin fibers (Jiang and Pugh 2009). Importantly, however, unlike nucleases, DNMT probing maintains the integrity of DNA strands, providing a single-molecule in addition to population-ensemble view of chromatin accessibility. Moreover, the single-molecule output identifies epigenetic variation across populations of cells and chromatin configurations unique to minority subpopulations of cells (Darst et al. 2013; Nabilsi et al. 2014), without the need for costly, specialized single-cell approaches (Jin et al. 2015; Lu et al. 2016; Lai et al. 2018;

Mezger et al. 2018). Whereas isolation of mammalian cell nuclei for MAPit is readily achieved by cell lysis with non-ionic detergents, adaptation to plant tissues requires additional disruption of the cell wall and removal of tissue debris. We adopted a method of tissue grinding with a mortar and pestle in nuclei isolation buffer (NIB) containing metal ion chelators and polyamines on ice to stabilize the structure of nuclei and native chromatin (**Supplemental Fig. 7**). When fluorescence-activated nuclei sorting (FANS) was employed, tissue was finely chopped as previously described (Lu et al. 2017). Nuclei from independent biological duplicates of *Arabidopsis thaliana* Col seedlings were collected through filtration and density centrifugation. The purified nuclei were separated into two aliquots and incubated in parallel with M.CviPI and enzyme diluent only (mock) exactly as done previously with mammalian cells (Pardo et al. 2011; Nabilsi et al. 2014; Pardo et al. 2015). Whole genome bisulfite sequencing or targeted sequencing of mock samples served to determine levels of native DNA methylation, which were subtracted from the M.CviPI-probed samples to reveal the net level of chromatin accessibility at each GC cytosine. In targeted analysis, two tricolor schemes were used to plot endogenous methylation and M.CviPI-dependent methylation.

**Genome-wide profiling of chromatin accessibility in plants**

The overall digestion level of chromatin with nucleases, including DNase I, MNase and Tn5 transposase, must be titrated to avoid over-digestion and hence false assignment of regions as accessible (Buenrostro et al. 2015; Chereji et al. 2019a; Liu et al. 2019). By contrast, MAPit is performed at or near saturating levels of exogenous DNMT and retains the continuity of epigenetic and genetic content of individual DNA strands, allowing for the application of single-molecule and -cell sequencing (Guo et al. 2017; Pott 2017; Clark et al. 2018; Li et al. 2018). The elimination of nuclease digestion allows a wider range of enzyme doses to be used in the probing reaction. A dose of 30-100 units of M.CviPI per million human cells have been used, while for human and mouse single cells, 2-5 units per cell have been successfully applied (Nabilsi et al. 2014; Pardo et al. 2015; Guo et al. 2017; Clark et al. 2018; Li et al. 2018).

Although MAPit has been thoroughly vetted in studies of mammalian cells, such a large genome has not been surveyed by a multi-point and hence costly titration of M.CviPI activity. This would add a quantitative dimension in that reaching a plateau delineates the fractions of a genome that are accessible to probe (Chereji et al. 2019b). To establish our MAPit workflow in *Arabidopsis thaliana*, a model plant with a relatively small 135 Mb genome, nuclei isolated from leaves of two independent sets of plants grown under each condition by cold grinding and FANS were incubated with increasing doses of M.CviPI. The resulting MAPit followed by methylome analysis, i.e., MAPit-WGBS, for each pair of replicates was highly reproducible (**Supplemental Fig. 8**). A plot of the genome-wide average GC methylation level *versus* M.CviPI units per million *A. thaliana* nuclei yielded a dose response curve that plateaued at 100U/$10^6$ nuclei, with ~55% overall $Gm^5C$ content (**Supplemental Fig. 9**). A dose of 100 U M.CviPI per million mammalian nuclei is also sufficient to achieve saturation of nucleosome-free regions (NFRs) of numerous analyzed promoters within diverse mammalian cell types (Pardo et al. 2011; Darst et al. 2013; Nabilsi et al. 2014; Pardo et al. 2015; Stees et al. 2016). The average $Gm^5C$ level of the fresh tissue, 22°C-100 U sample prepared from ground leaf tissue, i.e., without FANS, was close to the saturated level, (**Supplemental Fig. 9A**, dotted line). At the level of a representative chromosome (**Supplemental Fig. 10**), the levels of $Gm^5C$ increased with dose, mirroring **Supplemental Fig. 9A**. Furthermore, the region-to-region pattern of accessibility was strikingly similar over the range of employed enzyme doses (**Supplemental Fig. 9B** and **Supplemental Fig. 10**). These data suggest that accessibility to M.CviPI in nuclei isolated from *Arabidopsis* is region-specific and reaches stable, saturated maxima.

We performed a more granular meta-analysis of the MAPit-WGBS data focusing on sequences within 1 kb upstream and downstream of TSSs across the genome. All four doses of M.CviPI displayed preferential accessibility of TSSs compared with flanking regions of chromatin (**Supplemental Fig. 9B**). On closer examination, the patterns of promoter accessibility were remarkably similar for the 30 and 100 U/million nuclei. The latter dose was chosen for further experimentation because accessibility of nucleosome-depleted TSSs approached that of the highest dose, while maximizing differential accessibility between the TSSs and flanking regions of chromatin.

In mammalian cells where non-CG methylation is not detectable, M.CviPI and endogenous DNMTs can both methylate GCG, and therefore only HCG and GCH are examined (Pardo et al. 2011; Kelly et al. 2012). Although some resolution is sacrificed, unequivocal assignment exogenous versus endogenous methylation is achieved and a mock (0 U) sample can be bypassed. In plants, however, the abundance of non-CG methylation dictates a different approach. Specifically, a mock reaction is conducted in parallel to determine baseline $Gm^5C$, which is then subtracted from the level of in DNA obtained from M.CviPI-probed nuclei. The validity of this approach is supported by three observations: 1) high consistency between mock and probed samples in levels of $Hm^5C$, i.e., excluding $Gm^5C$ (**Supplemental Fig. 9C**), demonstrating that native levels of methylation are unaffected by introduction of $Gm^5C$; 2) on average, baseline $Gm^5C$ was low, ~8% genome-wide (**Supplemental Fig. 9A**, 0 U point) and < 8% within 1 kb upstream and downstream of TSSs (**Supplemental Fig. 9D**); 3) in general, background hypermethylated GC co-localized with hypo-accessible regions (**Supplemental Fig. 11**), such as was observed approaching 1 kb upstream of TSSs (**Supplemental Fig. 9D**). This is consistent with the reported negative correlation between all three types of DNA methylation and MNase accessibility in *A. thaliana* (Zhao et al. 2020). Therefore, when baseline, endogenous $Gm^5C$ is high, the difference between non-probed and probed samples should be small, reflecting a low accessibility. After subtraction of baseline $Gm^5C$, the 22°C-100 U sample showed a GC accessibility maxima centered around the TSSs (**Supplemental Fig. 9D** and **9E**), which was consistent with accessible regions identified in *A. thaliana* genes (Lu et al. 2017).

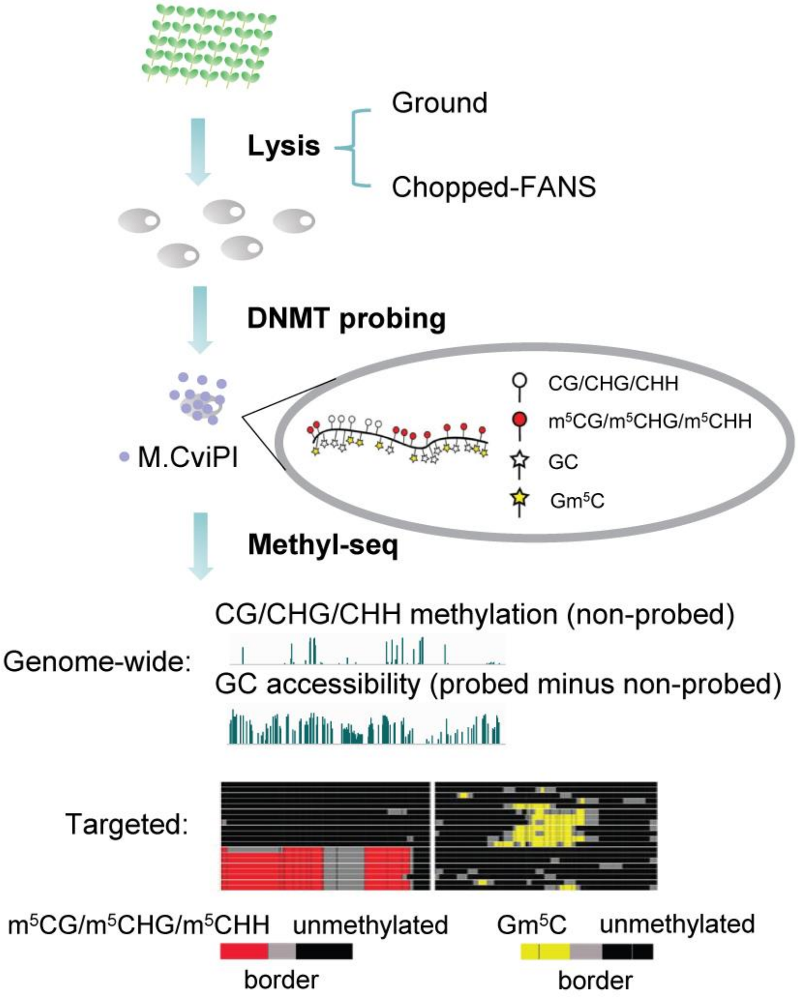

**Supplemental Figure 7.** MAPit workflow in plants. Plant tissues are ground in ice-cold nuclei isolation buffer using a pre-chilled mortar and pestle. Chopping fresh tissues is preferred when FANS is employed. The isolated nuclei are probed for accessible DNA in chromatin by an exogenously added DNMT, in this study, M.CviPI targeting GC dinucleotides. Purified DNA is subjected to WGBS or targeted methylation sequencing. Chromatin accessibility is assessed by subtracting the GC methylation levels (Gm$^5$C) of the mock-probed from the probed samples. In parallel, native DNA methylation at CG, CHG and CHH sites is also investigated in mock-probed samples. For targeted single-molecule level analysis, tri-color plot are used to represent chromatin accessibility and DNA methylation of individual molecules of chromatin within the original M.CviPI-treated *Arabidopsis* nuclei. The molecules are plotted in the same order along the y-axis. Yellow connects two or more consecutive G$m^5$C are with, whereas red connects two or more consecutive $m^5$CG, $m^5$CHG and $m^5$CHH in their respective panels. Black connects two or more consecutive non-methylated C, and gray connects the transitions between a non-methylated C and $m^5$C in any context as a border.

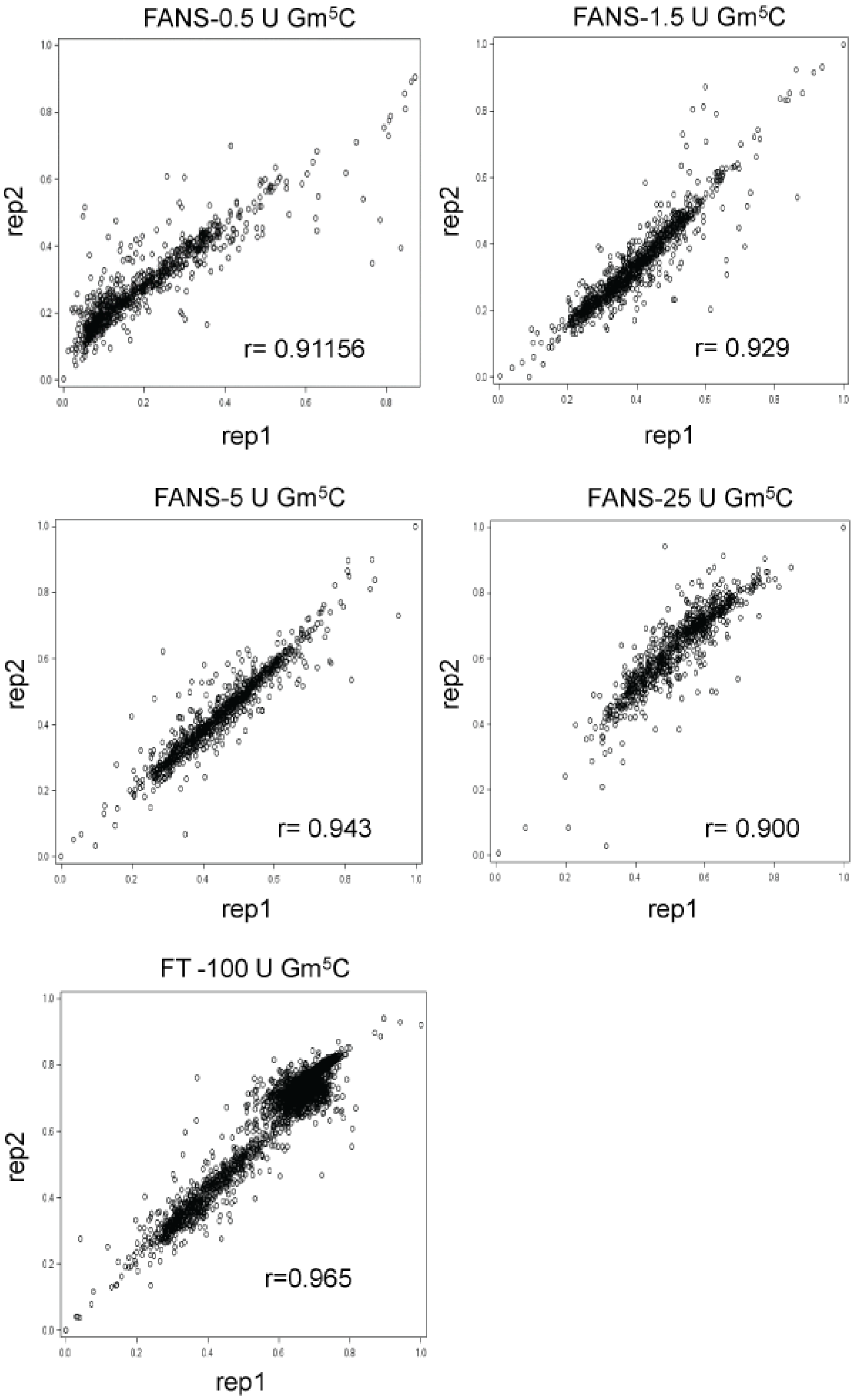

**Supplemental Figure 8.** Correlation of cytosine methylation levels between two biological replicates determined by MAPit-WGBS. Correlation of CG/CHG/CHH methylation levels (**A**) and GC methylation levels (**B**) are shown. (**C**) FANS samples are analyzed in GC context.

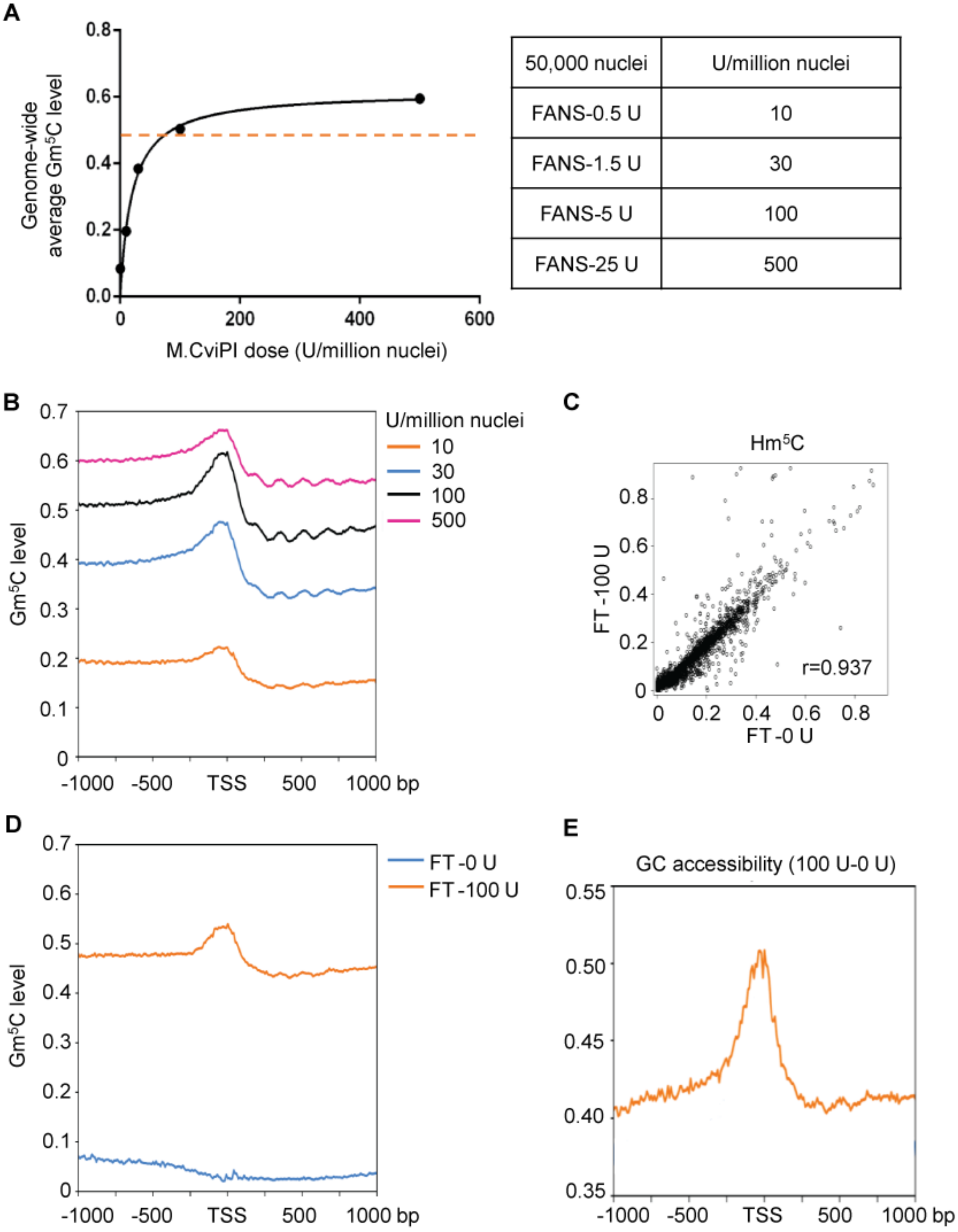

| 50,000 nuclei | U/million nuclei |
|---|---|
| FANS-0.5 U | 10 |
| FANS-1.5 U | 30 |
| FANS-5 U | 100 |
| FANS-25 U | 500 |

**Supplemental Figure 9.** Overview of genome-wide chromatin accessibility detected by MAPit-WGBS in *Arabidopsis*. (**A**) Genome-wide average $Gm^5C$ level of samples treated without (0 U) or with increasing doses of M.CviPI. The dose response was conducted on FANS samples obtained from plants grown at 22°C. The orange dotted line indicates the level of $Gm^5C$ in the 100 U sample obtained from ground tissue, i.e., without FANS. (**B**) Average $Gm^5C$ levels within –/+ 1 kb of the TSS plotted using 200-bp bins for FANS samples probed with the indicated doses of M.CviPI. (**C**) Correlation of native $Hm^5C$ levels between samples treated with 0 U and 100 U M.CviPI. (**D**) Average $Gm^5C$ levels are plotted in the same way as (**B**) for pooled samples. (**E**) Average chromatin accessibility –/+ 1 kb of the TSS are plotted using 200-bp bins for the pooled samples. Net accessibility was obtained by subtracting GC methylation levels of the 0 U samples from the 100 U samples. FANS=fluorescence-activated nuclei sorting,  FT=fresh tissue

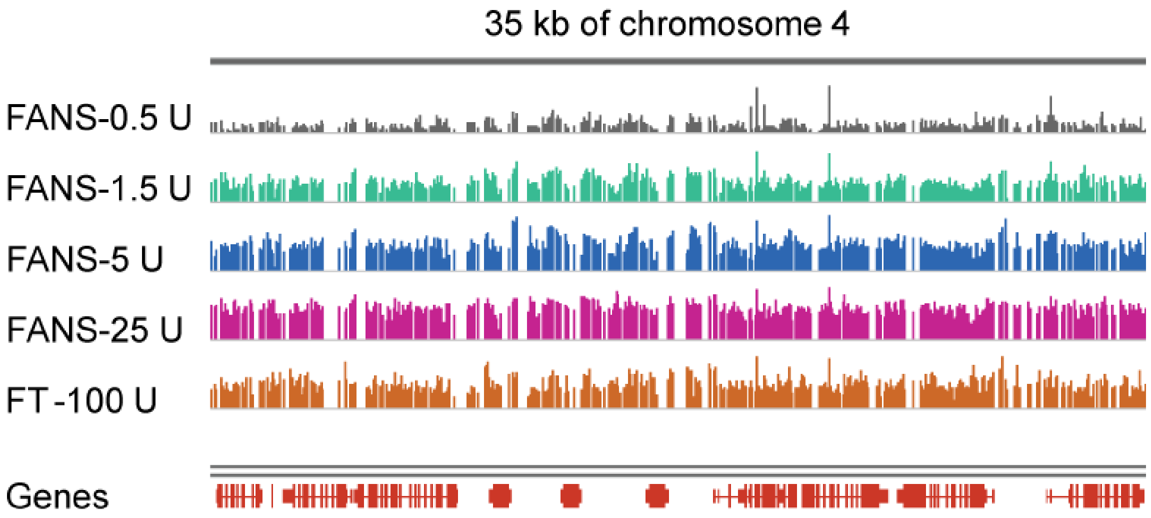

**Supplemental Figure 10.** Genome browser tracks of a representative 35 kb region from chromosome 4 comparing MAPit-WGBS at different doses of M.CviPI in *Arabidopsis* seedlings. The scale of all tracks is 0 to 1 showing Gm$^5$C level. FANS=fluorescence-activated nuclei sorting, FT=fresh tissue

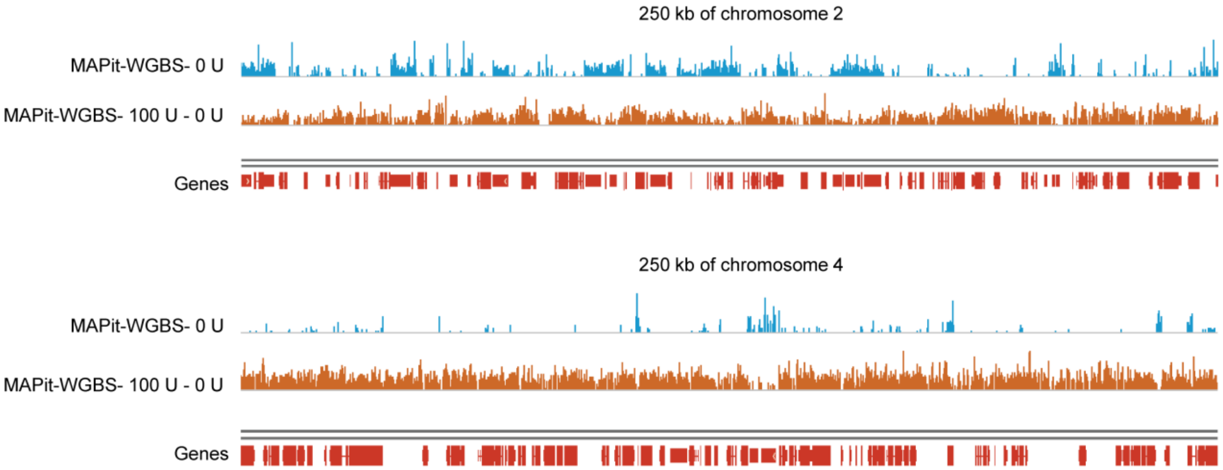

**Supplemental Figure 11.** Genome browser tracks of two representative 250 kb regions from chromosome 2 and 4 showing GC methylation levels and chromatin accessibility detected by MAPit-WGBS. The scale of all MAPit-WGBS tracks is 0 to 1.

**Supplemental Protocol.** MAPit for plant tissues

**Reagents**

Nuclei Isolation Buffer (NIB) for MAPit: 15 mM Tris, pH 7.8, 0.30 M sucrose, 0.3 mM spermine, 0.125mM spermidine, 20 mM NaCl, 80 mM KCl, 15 mM β-Mercaptoethanol, 0.1 mM EGTA, 1 mM EDTA, 1/100 volume of protease inhibitor cocktail (inhibiting aspartyl, cysteine, and serine proteases, as well as aminopeptidases), 0.15% Triton X-100.

Cell resuspension buffer (CRB): 20 mM HEPES pH7.5, 70 mM NaCl, 0.25 mM EDTA, 0.5 mM EGTA, 0.5% glycerol, 10 mM DTT (fresh add), 0.25 mM PMSF (fresh add).

Methylation buffer (MB): 320 µM *S*-adenosyl-*L*-methionine in CRB.

Methylation stop buffer (MSB): 100 mM NaCl, 10 mM EDTA, 1% SDS.

**Procedure**

1. Collect fresh plant tissues to cold (0-4°C) NIB. The amount and type of tissues need to be tested and adjusted in preliminary experiments. If tissues frozen in liquid nitrogen are used, thaw in the pre-chilled mortar on ice.
2. Grind tissues in a pre-chilled mortar and pestle on ice, with 5 ml of cold (0-4°C) NIB.
3. Gently add cold NIB using plastic pipette to bring the final volume to 10 ml and homogenize it. Remove cell debris from the nuclear suspension by filtration though a four-layer bundle of cheesecloth, one layer of miracloth, and one 105-µm mesh polypropylene screen. All steps should be done on ice.
4. Pellet nuclei by centrifugation at 1500g at 4°C for 10 min. Discard supernatant.

5. Gently add 2-3 ml cold NIB into the tube using plastic pipette. Do not break the pellet. Use a small brush to gently resuspend the nuclei in NIB, leaving white starch in the pellet.
6. Transfer the slurry to a new tube using plastic pipette. Add cold NIB to 5 ml and re-spin at 1500g or 3000 rpm at 4°C for 10 min. Discard supernatant. Step 5 and 6 can be omitted if young tissues are used.
7. Add 200 µl CRB to wash the pellet, respin. Discard supernatant.
8. Add 200 µl MB to resuspend the nuclei. The volume of methyl buffer can be adjusted depending on how many reactions will be needed.
9. Pre-warm nuclei in 37°C for 2 min. Pre warm MSB in 70°C for 10 min.
10. Aliquot 98 µl of nuclei and add 2 µl methyl buffer (0 U) or M.CviPI (50 U/µl stock) (100 U). Incubate in 37°C for 15 min. The dose if M.CviPI should be adjusted according to the tissue amount and genome size.
11. Add 100 µl pre-warmed MSB and mix well.
12. Add 2 µl RNase A (1 mg/ml) (final 10 ug/ml) in RT for 1 h.
13. Add 1 µl Proteinase K (20 mg/ml) (final 100 ug/ml) and incubate in 50°C overnight.
14. Add 200 µl Phenol/chloroform/isoamyl alcohol (25:24:1) and vortex.
15. Spin 10 min at RT and transfer aqueous phase (upper) to new tube (around 162 uL).
16. Add 54 µl 10 M AmOAc (final 2.5 M) (mix gently by snapping with finger) and 500 µL 100% EtOH (mix gently by inverting tube a few times – do not vortex). Put at -20°C at least 2 hours.
17. Spin at 4°C for 10 min. Decant carefully and then wash pellet with 70% EtOH to help get rid of residual AmOAc (can spin again if pellet becomes dislodged).

18. Dry upside down on Kimwipes. Resuspend pellet in 50 uL 0.1 x TE or $ddH_2O$ (depending on needs/pellet size).
19. The saturation of GC methylation can be tested through digestion using HaeIII, a methylation sensitive enzyme targeting GGCC, followed by quantitative PCR on selected region, or targeted methylation sequencing on samples with gradient doses of M.CviPI.

**Supplemental Note 2.** Epigenetic changes proximal to transposable elements in *Arabidopsis* in the response to ionizing radiation

CHH methylation is known to be enriched near and may regulate the expression of transposable elements (TEs) (e.g., (Martin et al. 2021)). TEs can be activated in response to biotic and abiotic stress (such as ionizing radiation (IR)), yet owing to their repetitive nature, are difficult to quantify transcriptionally due the inability to accurately assign sequence reads to specific elements. However, differential methylation of CHH regions associated with TEs, if any, could allow us to infer if transposons are likely activated or repressed in response to IR. We annotated CHH DMRs to transposons and compared their annotated feature (1 kb-upstream, transposon body, 1 kb-downstream) to their annotated genic feature (promoter, exon, intron, downstream, intergenic). We found that the majority of CHG DMRs (85% - 93%) and CHH DMRs (78% - 89%) were annotated to transposons, and most of those exclusively annotated to a transposon (CHG DMRs: 73 - 79%; CHH DMRs: 62 - 66%; **Supplemental Table 7**). Between 11 - 18% of CHG DMRs and 16 - 23% of CHG DMRs were located close to both a protein-coding gene and a transposable element (**Supplemental Table 7**). In contrast, the proximity of CG DMRs to genes and TEs was more dose-dependent, although about 20% of all CG DMR were located near to both a gene and a TE (**Supplemental Table 7**). Examining chromatin accessibility and endogenous methylation across TE bodies (± 1kb) revealed reduced accessibility and increased methylation at TE bodies was lower than sequences up- and downstream of the TE (**Supplemental Fig. 12**). Accessibility at and near TEs was generally higher after IR exposure, following a similar trend to accessibility at/near genes (**Supplemental Fig. 12A** and **12B**). CG and CHG methylation were lower exclusively for 10 cGy compared to Mock (**Supplemental Fig. 12B** and **12C**), while TE methylation for 100 cGy was unchanged. In contrast, CHH methylation at TEs displayed dose-dependent changes, with decreased CHH methylation for 10 cGy but increased methylation for 100 cGy compared to Mock (**Supplemental Fig. 12D**).

We queried if DMRs annotated to TEs were more likely to also overlap at least one DAR. A statistically significant increase in proportion of TE-annotated DMRs overlapping DARs compared to non-TE-annotated DMRs was observed for CHH DMRs (10 cGy and 100 cGy IR exposure), but less consistently for the other methylation types (**Supplemental Table 8**). While the difference is small, this suggests that while altered TE regulation may be consequence of IR exposure it is not likely a main component of response to IR.

**Supplemental Table 7.** Genic and transposable element annotation of differentially methylated regions

File: Supplemental_Table_7_TE_vs_Gene_annotation_DMRs.xlsx

**Supplemental Table 8.** Comparison of differentially accessible regions and transposable elements within differentially methylated regions

File: Supplemental_Table_8_TE_DARs_DMRs.xlsx

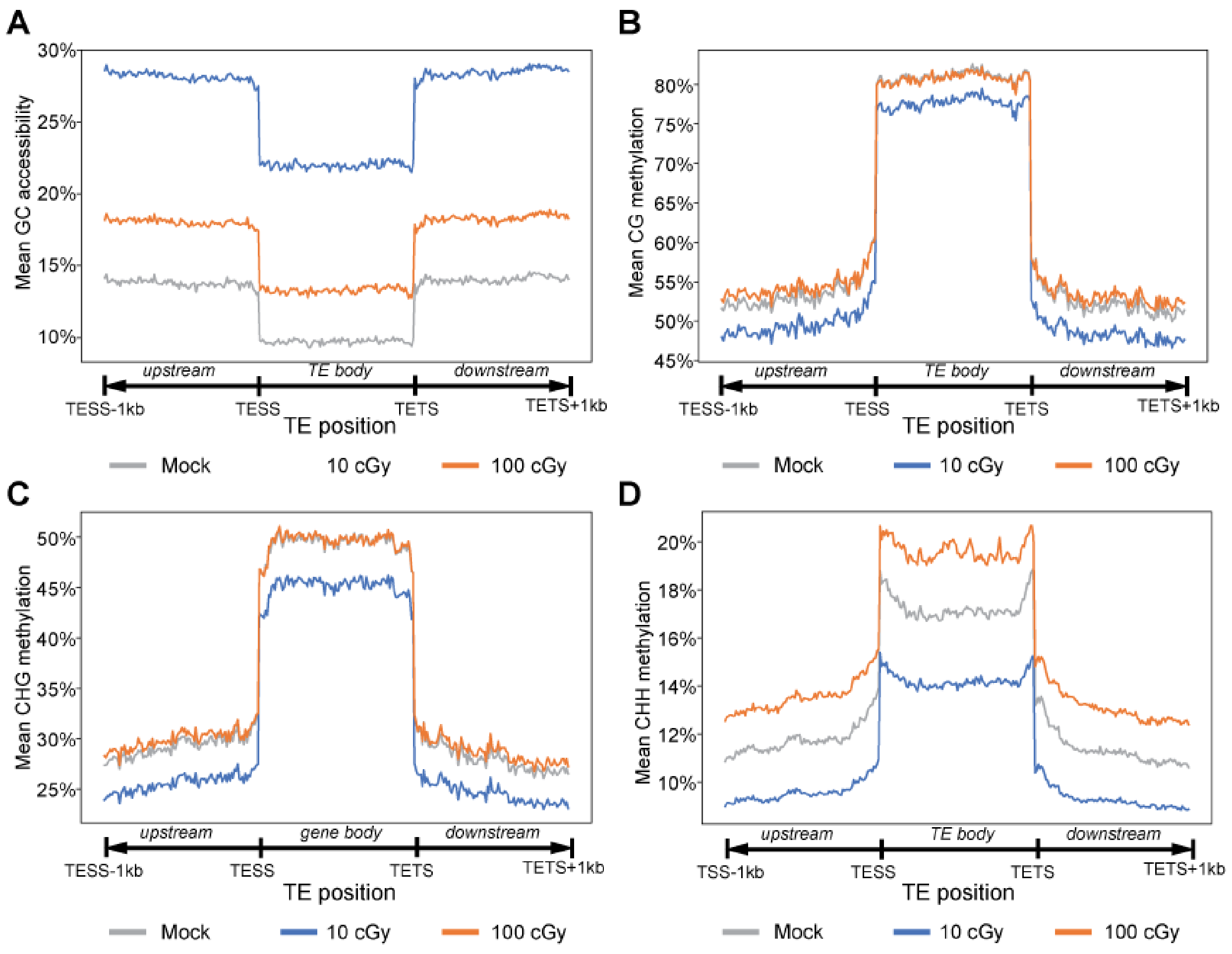

**Supplemental Figure 12.** Average methylation and accessibility across transposable element bodies. (**A**) Average GC accessibility across transposable element (TE) body (± 1kb) for each ionizing radiation (IR) exposure (Mock (grey), 10 cGy (blue), 100 cGy (orange)). Average (**B**) CG methylation, (**C**) CHG methylation, and (**D**) CHH methylation across TE regions (1kb 5' upstream, TE body, 1kb 3' downstream) for each IR exposure (Mock (grey), 10 cGy (blue), 100 cGy (orange)). TE body lengths were rescaled to 1kb. TESS = transposable element start site; TETS = transposable element termination site.

**Supplemental Note 3.** Differential methylation and accessibility model selection

Choice of an appropriate algorithm or model must be based on the limitations of the experimental design and how the response variable is measured or estimated. For the latter in whole genome studies of methylation, the experimental response can be measured as either (1) integer counts at each site of methylation ($m^5CG$, $m^5CHG$, $m^5CHH$, $Gm^5C$, or any other genomic sequence motif that is targeted by methylating/demethylating enzymes) as the number of reads corresponding to methylated cytosines and the number of reads corresponding to unmethylated cytosines; or (2) an estimate of proportion of methylated cytosines for a given site of methylation.

Accurate estimation of methylation frequency in an experimental condition is dependent on the sequencing coverage at a given genomic methylation site as well as overall sample size: methylation frequency can be over/underestimated at sites of low coverage can interfere with detection of differentially methylated sites/cytosines (DMCs) leading to increased variance and decreased statistical power. Conversely, very high coverage can amplify small, biologically irrelevant differences if estimates of variances are too small.

For our MAPit data for examining changes in methylation and chromatin accessibility in response to exposure to ionizing radiation (IR), we therefore tested several algorithms and models to determine the optimal analytical approach to our study.

**Dataset**

The distribution of coverage for all 42,329,792 called methylation sites from the MAPit data used in this study was 16.0 ± 133.6 reads (interquartile range 4 to 16 reads), and 31.0 ± 261.8 reads (interquartile range 8 to 31 reads) when counts from both biological replicates are summed.

We removed sites of low coverage (<10 total reads summed across both replicates per condition) and sites that were not called for both Mock-exposed and either 10 cGy or 100 cGy; the distribution of coverage for these 22,614,251 sites was 38.9 ± 113.6 reads (interquartile range 19 to 45 reads), suggesting this filtered dataset set had adequate coverage for subsequent analysis. Coverage between replicates (Pearson R = 0.97 – 1.00) and conditions (Pearson R = 0.97 – 1.00)  was well-correlated, suggesting that imbalances in coverage between conditions would not greatly impact our findings. However, the experimental design consists of only 2 replicates per condition. To test for differential methylation, we therefore require a statistical test that is appropriate for small sample sizes while also sensitive to differences in overall coverage. We therefore explored several models including statistical software developed for the analysis of methylation data to identify with method would most appropriate model our data.

**Model selection**

We then selected several published analysis software and other approaches to use for determining which analytical approach would work best for our data. While there are many options for simple two-group DMC/DMR detection there are few tools that can be extended to include covariate data, which would be required to assess changes in Gm$^5$C accessibility while controlling for endogenous methylation levels. Therefore, we had to exclude several well-used programs from our analysis as these algorithms could not be used to account for differences in Gm$^5$C methylation when accounting for the addition/absence of the M.CviPI enzyme. The algorithms selected for testing are summarized in **Supplemental Table 9**. For each of the selected models, replicates were treated as independent samples except where noted, and the comparisons 10 cGy vs Mock and 100 cGy vs Mock were analyzed separately.

The first approach does not utilize statistical analysis, and differentially methylated cytosines (DMCs) simply refers to any site with a difference in methylation above/below some fixed threshold (e.g. all $m^5$CG sites with a difference in methylation > 10% or < -10%, IR-exposed vs mock-exposed. For the present study the difference in methylation is calculated as % methylation for IR-exposed condition - % methylation for Mock-exposed condition, where % methylation is the proportion of cytosines that are methylation at a given site). As this is just a simple difference calculation, it can be readily adapted for defining differentially accessible cytosines (DACs) as any site with a difference in G$m^5$C methylation after addition of the M.CviPI enzyme correcting for endogenous methylation as: difference in % accessibility = [(% GC methylation for IR-exposed with 100 U M.CviPI - % G$m^5$C methylation for IR-exposed with 0 U M.CviPI) - (% G$m^5$C for Mock-exposed with 100 U M.CviPI - % G$m^5$C for Mock-exposed with 0 U M.CviPI)], where % G$m^5$C is the proportion of cytosines that are methylation at a given G$m^5$C site.

The second algorithm is Dispersion Shrinkage for Sequencing (DSS) a Bioconductor package for comparing differences in methylation profiles different conditions (Park and Wu 2016). DSS models methylation data as a beta-binomial distribution and estimates dispersion. It tests differential methylation using a Wald test on counts of methylated and unmethylated cytosines at each methylation site. DSS also allows for the use of covariates and building of custom models so it can be readily adapted for testing differential accessibility.

The third software tested was methylKit (Akalin et al. 2012), an R package that is used to model DNA methylation. It uses one of two approaches depending on sample size, both of which use methylated/unmethylated counts as input. When there is one sample per group, methylKit defaults to a Fisher's exact test (FET). When there are multiple samples per group, the data is assumed to be distributed binomially and the log-odds ratio of methylation proportion of a site or region is modeled logit($\pi_i$) = $\beta_0 + \beta_1 d_i$, where $d_i$ is experimental treatment (i.e. IR exposure). The regression approach also allows for the use of covariates, so it can also be applied to testing differential accessibility. The log-odds ratio of accessibility proportion of a site or region can therefore be modelled as logit($\pi_{ii}$) = $\beta_0 + \beta_1 d_i + \beta_2 m_j + \beta_3 d_i m_j$, where $d_i$ is experimental treatment (i.e. IR exposure) and $m_j$ is M.CviPI units. Methylkit also calculates a $Q$ statistic to correct for multiple hypothesis testing using a sliding linear model (Wang et al. 2011).

The fourth program tested was methylSig (Park et al. 2014), which can model counts of methylated and unmethylated cytosines using either a binomial regression or a beta-binomial regression. The binomial regression is similar to that implemented in methylKit. However neither the implemented binomial or betabinomial regressions allow for covariates in its model, and thus only a single predictor can be used. We therefore used methylSig to compare the output of its two regression approaches as a benchmark for only the analysis of endogenous methylation.

We also employed the Mann-Whitney U test as implemented in metilene (Jühling et al. 2016). Unlike DSS, methylKit, and methylSig, metilene uses % methylation only, thus any differences in coverage will be ignored. While metilene does not allow for covariates, for assessing differences in accessibility the % $Gm^5C$ accessibility can be calculated instead of % $m^5C$ for each combination of IR dose × M.CviPI units and modelled using the Mann-Whitney U test.

We also several model options as implemented in SAS (version 9.4, SAS Institute). The first of these was the use of FET, which we used to test differential endogenous methylation between IR- and Mock-exposed plants. Because FET does not allow for the adjustment of other covariates, for GC methylation it can only be employed to test if a $Gm^5C$ site within an exposure was differentially methylated between 0 U and 100 U M.CviPI, where a significant increase in GC methylation with M.CviPI corresponds to a site of open chromatin (as the M.CviPI enzyme is able chemically modify the cytosine). To account for the expected difference in GC methylation with the addition of M.CviPI, a Cochran-Mantel-Haenszel (CHM) test was used to assess differences in GC methylation between IR-exposed and Mock-exposed controlling for the addition of the GC methyltransferase. A possible limitation of the CMH test is that it does rely on the assumption that effect of treatment is relatively homogenous in all strata (i.e. with and without M.CviPI), and therefore if the difference in endogenous Gm$^5$C methylation between two conditions at a given site is directionally different from that of the difference in M.CviPI-treated Gm$^5$C methylation then these may be inadvertently missed.

We also applied three regression-based approaches, namely binary logistic regression, binomial logistic regression, and linear mixed models. Both binary logistic regression and binomial logistic regression will give the same outcome for testing DMC and DAC as binary logistic regression is a special case of the binomial logistic regression where the dependent variable has only two levels. In the case of our study, this is whether or not a sequencing read that maps to a specific locus corresponds to a methylated cytosine or an unmethylated cytosine. For linear mixed models, % m$^5$C methylation or % Gm$^5$C accessibility is modeled against predictor variables.

**Site-based analysis**

For site-based analysis of endogenous methylation, we used FET, binomial logistic regression, binary logistic regression, DSS, methylKit, methylSig, metilene, and "credible methylation difference" to identify DMCs. For logistic regression, we modelled methylation as: $y_i \sim B(r_i, \pi_i)$, where $y_i$ is the number of methylated reads for site $i$, distributed binomially with parameters $r_i$ and $\pi$; $r_i$ is the number of mapped reads for site $i$, and $\pi$ is the latent probability estimated logit($\pi_i$) ~ $\beta_0 + \beta_1 d_{ij}$; $\beta_0$ is a fixed intercept term and $d$ is IR dose (Mock vs 10 cGy or 100 cGy). The models used by methylkit, methylsig, DSS, and metilene were all run with only IR dose as the predictor term; for metilene, the % methylation was used as the response variable, while methylated/unmethylated cytosine count was used for all other applications.

As using credible methylation difference would result in an overabundance of potential DMCs (**Supplemental Fig. 13A**), we opted to use methylation difference in conjunction with one of the above statistical models. We counted the number of $m^5$CG, $m^5$CHG, and $m^5$CHH sites with differences in methylation between IR-exposed and Mock using criteria of ≥ 10%, ≥ 20%, ≥ 30%, or ≥ 50%, to determine what the minimum credible difference in methylation would be optimal (**Supplemental Fig. 13A**). As expected, the number of sites decreased with the minimum methylation difference increased (**Supplemental Fig. 13A**). We find that only 13% of the total analyzed methylation sites had a difference in methylation between IR-exposed and Mock of at least 10%, suggesting that changes in methylation in response to IR exposure are generally small.

We then applied each of the statistical models above to endogenous methylation data, analyzing each site type ($m^5$CG, $m^5$CHG, $m^5$CHH) and IR exposure (10 cGy vs Mock, 100 cGy vs Mock) separately. Metilene and methylKit internally correct for multiple testing by calculating a $Q$ value (ref?); for all other models, a False Discovery Rate (FDR) correction was applied. Statistical significance was defined as a $Q < 0.05$ or FDR-corrected $P < 0.05$. For SAS-implemented FET and logistic regression models, we also examined DMCs with an FDR-corrected $P < 0.01$ to examine the effect of significance threshold on number of DMCs. We found that the binomial regression implemented in methylSig produced overall highest number of DMCs, although this sharply decreased when a minimum methylation difference criterion was applied

(**Supplemental Fig. 13B**). DSS found 10 DMCs for 10 cGy but found substantially more for 100 cGy, although most of these involved sites with large differences (> 50%) in methylation between IR-exposed and Mock, metilene and the betabinomial regression test in methylSig found no significant DMCs. The best performing analysis methods were FET and the logistic regression approaches, which gave largely similar sets of DMCs (**Supplemental Fig. 13B**). As expected, imposing a stricter significance threshold (FDR $P < 0.05$ to FDR $P < 0.01$) decreased the overall number of DMCs found using FET or logistic regression but not greatly. MethylKit also performed well and also found most of the same DMCs, although fewer overall DMCs were detected with this approach (**Supplemental Fig. 13B**).

Given the small sample size (N = 2 per condition) and the relatively few individual DMCs called overall, for the purposes of our study we defined a DMC as site of endogenous methylation ($m^5CG$, $m^5CHG$, $m^5CHH$) with FET FDR-corrected $P < 0.05$ and a minimum methylation difference of 10%. This maximizes the number of possible DMCs and in addition many of these are also significant using the other analysis methods. Given the number of statistically significant DMCs (FDR $P < 0.05$), increasing the minimum methylation difference criterion too high would result in a potentially large loss of information, and thus we opted to use 10% as the minimum credible difference in methylation between conditions for subsequent analysis to maximize our detection of potentially biologically meaningful DMCs.

As plants have more types of DNA methylation than do typical mammalian systems ($m^5CG$, $m^5CHG$, $m^5CHH$ vs $m^5CG$), and 89% of $Gm^5C$ sites overlap with one of these sites of endogenous methylation (**Supplemental Fig. 13C**), model selection is critical for using $Gm^5C$ for determining chromatin accessibility from $Gm^5C$ methylation. For site-based analysis of chromatin accessibility, assayed through GC methylation induced by M.CviPI, we used CMH test, binomial logistic regression, binary logistic regression, DSS, methylKit, metilene, and "credible accessibility difference" to identify DACs. FET was applied to assess differences in $Gm^5C$ methylation with and without the addition of the M.CviPI enzyme to identify $Gm^5C$ sites with M.CviPI-induced methylation significantly above background methylation, but as this is limited to a two-dimensional crosstabulation (i.e. a 2×2 table of methylation status vs IR dose or methylation status vs M.CviPI units) cannot used for testing if differences in $Gm^5C$ methylation in this experiment as that requires three dimensions (i.e. 2×2×2 table of methylation status, IR dose, and M.CviPI units). Instead, we applied the closest equivalent test, the CMH test, to assess differences in $Gm^5C$ methylation between IR exposures and used M.CviPI units (0 U, 100 U) as the stratifying variable. For logistic regression, we modelled methylation as: $y_i \sim B(r_i, \pi_i)$, where $y_i$ is the number of methylated reads for site $i$, distributed binomially with parameters $r_i$ and $\pi$; $r_i$ is the number of mapped reads for site $i$, and $\pi$ is the latent probability estimated $\text{logit}(\pi_i) \sim \beta_0 + \beta_1 d_{ij} + \beta_2 m_{ij} + \beta_3 d_{ij} m_{ij}$; $\beta_0$ is a fixed intercept term, $d$ is IR dose (Mock vs 10 cGy or 100 cGy) and $m$ is M.CviPI units (0 U, 100 U). The interaction term between IR dose and M.CviPI units, $d_{ij} m_{ij}$, was used as the predictor variable for differential accessibility. The models used by Methylkit and DSS were run with IR exposure (Mock vs 10 cGy or 100 cGy), M.Cvi units (0 U, 100 U) and the interaction IR exposure × M.Cvi units as the predictor terms; for Metilene, the % accessibility was used as the response variable, and a simple two-group comparison of IR exposure (Mock vs 10 cGy or 100 cGy) was performed. As with the DMC analysis, for Metilene and MethylKit, statistically significant DACs were defined as having a $Q < 0.05$; FDR correction was applied for all other DAC tests and statistical significance was defined FDR-corrected $P < 0.05$. For FET and logistic regression models implemented in SAS, we also examined DACs with an FDR-corrected $P < 0.01$ to examine the effect of significance threshold on number of DACs.

To determine an optimal minimum credible difference in accessibility, we counted the number of GC sites with differences between IR-exposed and Mock using criteria of ≥ 10%, ≥ 20%, ≥ 30%, or ≥ 50% difference in accessibility (**Supplemental Fig. 13D**). Similar to using a credible methylation difference as the sole criterion for calling differential methylation, using only the analogous credible accessibility difference to identify DACs resulted in ~60% $Gm^5C$ sites being called as DAC. A similar result was obtained when using only those sites with significant differences (FET FDR $P < 0.05$) in $Gm^5C$ methylation between 0 U and 100 U of M.CviPI for at least one IR exposure (**Supplemental Fig. 13D**). We therefore decided used the difference in accessibility in conjunction with a statistical test to identify DACs, as was done for DMCs.

No DACs were identified using binomial logistic regression nor Metilene (**Supplemental Fig. 13E** and **13F**). Other models were less consistent with calling DACs. Methylkit and DSS models both identified DACs only for 10 cGy IR exposure, though with vastly different results. Methylkit identified 5 DACs, while 61% of all $Gm^5C$ sites analyzed were considered differentially accessible using the DSS model (**Supplemental Fig. 13E**). The CMH test produced more consistent results for both IR exposures (1,052 DACs for 10 cGy, 993 DACs for 100 cGy; **Supplemental Fig. 13F**). The CMH test was thus selected as the statistical test for DACs, in part because of its greater consistency but also as its 2×2 analogue, FET, was chosen for testing DMCs. In terms of a minimum difference in accessibility as the second criteria, we noted a considerable decrease in DACs when using a minimum change in accessibility of 10% (**Supplemental Fig. 13F**). For consistency with the analysis of differential methylation, a DAC was defined as site of $Gm^5C$ methylation with CMH FDR-corrected $P < 0.05$ between methylation status and IR exposure, stratified for M.CviPI units, and with a minimum accessibility difference of 10%.

### Region-based differential methylation analysis

For each assayed endogenous methylation type ($m^5CG$, $m^5CHG$, and $m^5CHH$), methylation sites were grouped into regions if they were ≤ 100 bp apart and the difference in methylation between exposed and Mock-exposed was in the same direction, as described in the main manuscript (**Supplemental Fig. 6**). We did not use predefined methylation regions from other sources and instead opted for a data-driven approach to segment the genome into regions that may be under the same epigenetic regulation under the tested experimental conditions. We also did not use a tiling window strategy, as we expect region size to vary. For each region defined using out methylation site grouping strategy, we set limits on the minimum number of methylation sites comprising each region. We explored the number of regions that could be constructed with different minimum numbers of sites (2, 3, 5, or 10 sites) with a minimum difference in methylation of 10% or 20% (**Supplemental Fig. 14A** and **14B**). A minimum of 10 sites per region results in very few regions that can be tested for differential methylation, whereas too few sites would bias DMR length towards more, shorter regions and may adversely impact the false discovery rate. We therefore explored the impact of number of sites per region on DMR identification.

We then applied two strategies to DMR identification. The first method is DMC counting and setting a minimum number of DMC (FET FDR $P < 0.05$ and difference in methylation ≥ 10% or ≤ -10%) within a region as a criterion for calling DMRs in this manner. For this we explored using several minimum DMC counts (1, 2, 3, 5, or 10 DMCs) to identify DMRs. This is a similar approach to how DSS calls DMRs, as DSS tests for differential methylation at each individual site, then report regions with DMCs (Park and Wu 2016). However, we excluded DSS from DMR analyses as there were few DMCs in our data that were called using DSS and therefore would unlikely detected significant DMRs.

As expected, when region size is small (2 sites per region) and only a single DMC with an absolute methylation difference ≥ 10% between conditions the number of regions called as DMRs was the highest. The major determinant on number of DMRs called using this first, DMC-based approach is the minimum number of DMCs required for a region to be considered differentially methylated (**Supplemental Fig. 14A** and **14B**). Region size (as number of cytosines tested within the region) also affected the number of DMRs but less consistently on different methylation types, with $m^5$CG methylation more impacted by region size than $m^5$CHG or $m^5$CHH methylation (**Supplemental Fig. 14A** and **14B**).

The second method for identifying DMRs, which assesses if there are overall regional differences in methylation, was performed using the algorithms in **Supplemental Table 9** to test if regions of increased or decreased methylation were statistically significantly different between IR and Mock exposures. For methylKit, methylSig, and metilene we provided input data from both replicates of each condition. Both methylkit and methylSig summarize methylation information for all sites within a region and performs a logistic regression on the combined data to test for differential methylation. The betabinomial regression employed by methylSig also uses sample-specific methylation counts to take into consideration the variability between samples being compared (ref). For the binomial regression implemented in SAS, we modelled methylation in each region using each site within a region as a form of pseudoreplication. Using the GLIMMIX procedure methylation was modelled as: $y_i \sim \mathrm{B}(r_i, \pi_i)$, where $y_i$ is the number of methylated reads for site $i$, distributed binomially with parameters $r_i$ and $\pi$; $r_i$ is the number of mapped reads for site $i$, and $\pi$ is the latent probability estimated $\mathrm{logit}(\pi_i) \sim \beta_0 + \beta_1 d_j + u_i$; $\beta_0$ is a fixed intercept term and $d$ is IR dose (Mock vs 10 cGy or 100 cGy) and $u$ is a random intercept term for site $i$ and distribution $\sim N(0, \sigma)$. For the linear regression model methylation at each site within each region was used as pseudoreplication. We used the MIXED procedure and modelled methylation as: $y_{ij} \sim \eta_0 + d_i + \varepsilon_{ij}$, where $y_{ij}$ is % methylation for condition $i$ and site $j$, $\eta_0$ is a fixed intercept term, $d_i$ is IR dose (Mock vs 10 cGy or 100 cGy) and was the main predictor term, and $\varepsilon_{ij}$ is the residual. All tests were corrected for multiple comparisons either calculating a $Q$ statistic (methylKit) or FDR correction (methylSig, metilene, logistic regression, linear regression).

The outcome of model-based approaches was highly variable (**Supplemental Fig. 14C** and **14D**). The binomial and betabionomial regression tests used by methylSig yielded identical results and were grouped together for comparison with other models. Linear mixed modelling identified the overall most DMRs, however most of these were when regions consisting of two methylation sites or more were considered, and the number of significant DMRs dropped off as region size increased. The overall highest numbers of DMRs called, regardless of model, tended to be when region size was at least three or at least five sites. When a minimum region size of three methylation size was considered linear modelling and then methylKit identified the most DMRs regardless of minimum % methylation cut-off (**Supplemental Fig. 14C** and **14D**). On the other hand, when a minimum of 5 sites is considered, the binomial logistic regression model, linear mixed model, methylKit, and methylSig each identify comparable numbers of DMRs (**Supplemental Fig. 14C** and **14D**). We also that DMRs identified with metilene and methylKit are frequently also found to be statistically significant with the binomial regression model, suggesting that these methods identify largely the same set of DMRs.

Model-based DMR tests find more DMRs than simply counting regions with significant DMCs in our data, and in most cases the model-based analysis identify all significant regions identified with the DMC-counting method. For model-based DMR analyses, we decided that using a region size of at least 5 methylation sites or more would provide the most informative set. Fewer sites produced many more regions to test with fewer possible datapoints, while requiring more sites results in relatively few testable regions.

#### Region-based differential accessibility analysis

Regions for testing differential accessibility using background-corrected M.CviPI-induced GC methylation were defined as illustrated in **Supplemental Fig. 6**). Like analysis of endogenous methylation changes, e explored the number of regions that could be defined with different minimum numbers of sites (2, 3, 5, or 10 GC sites) with a minimum difference in accessibility of 10% or 20% (**Supplemental Fig. 15A** and **15B**). We counted DACs within each regions and examined setting a minimum number of DACs (CMH FDR $P < 0.05$ and difference in methylation ≥ 10% or ≤ -10%) within a region as a criterion for calling a DARs (minimum DAC counts: 1, 2, 3, 5, or 10 DACs). Few DARs were called in this manner for 100 cGy (**Supplemental Fig. 15A** and **15B**). By comparison, there were 5,627 regions with at least one significant DAR (10% difference in methylation) for 10 cGy, and generally had differences in accessibility >20%. However, most regions with DACs tended to be in smaller regions (consisting of < 5 GC sites) with only a single significant DAC present; few regions had 5 or more DACs present (**Supplemental Fig. 15A** and **15B**).

We then used the same algorithms that were used in model-based DMR analysis (**Supplemental Table 9**), to assess if there are overall regional differences in Gm$^5$C accessibility between IR-exposed and Mock-exposed plants. As with the DAC analysis, we excluded methylSig as it does not incorporate more than one predictor term. Analysis with methylKit summarized methylation information for all sites within a region for each condition (Mock 0 U M.CviPI, IR-exposed 0 U M.CviPI, Mock 100 U M.CviPI, IR-exposed 100 U M.CviPI) and a logistic regression on the combined data was performed. For the binomial regression implemented in SAS, we modelled Gm$^5$C accessibility in each region using each site within a region as a form of pseudoreplication. Using the GLIMMIX procedure methylation was modelled as: $y_i \sim B(r_i, \pi_i)$, where $y_i$ is the number of methylated reads for site $i$, distributed binomially with parameters $r_i$ and $\pi$; $r_i$ is the number of mapped reads for site $i$, and $\pi$ is the latent probability estimated $\text{logit}(\pi_{ijk}) \sim \beta_0 + \beta_1 d_j + \beta_2 m_k + \beta_1 d_j m_k + u_i$; $\beta_0$ is a fixed intercept term and $d$ is IR dose (Mock vs 10 cGy or 100 cGy) and $m$ is M.CviPI units (0 U, 100 U). The interaction term between IR dose and M.CviPI units was used as the predictor variable for differential accessibility; $u$ is a random intercept term for site $i$ and distribution $\sim N(0, \sigma)$.

Two linear mixed models were tested, with both using the methylation at each Gm$^5$C site within each region as pseudoreplication. The first linear mixed model used the MIXED procedure and modelled accessibility as: $y_{ijk} \sim \eta_0 + d_i + m_j + d_i m_j + \varepsilon_{ijk}$, where $y_{ijk}$ is % methylation at site $k$, $\eta_0$ is a fixed intercept term, $d_i$ is IR dose (Mock vs 10 cGy or 100 cGy), $m_j$ is M.CviPI units (0 U, 100 U), $d_i m_j$ is the interaction between IR dose and M.CviPI units and the main predictor term, and $\varepsilon_{ij}$ is the residual. The second linear mixed model used the MIXED procedure and modelled accessibility as: $y_{ijk} \sim \eta_0 + d_i + \varepsilon_{ijk}$, where $y_{ijk}$ is % Gm$^5$C accessibility (calculated as % methylation at 100 U M.CviPI - % methylation at 0 U M.CviPI) at site $k$, $\eta_0$ is a fixed intercept term, $d_i$ is IR dose (Mock vs 10 cGy or 100 cGy) and the main predictor term, and $\varepsilon_{ij}$ is the residual.

All tests were corrected for multiple comparisons either calculating a $Q$ statistic (methylKit) or FDR correction (metilene, logistic regression, linear regression).

Similar to DMRs, linear mixed modelling produced more statistically significant DARs, although most of these tended to be regions with fewer than five Gm$^5$C sites (**Supplemental Fig.**

**15C** and **15D**). Modelling % accessibility as a function of IR exposure yielded more significant regions than modelling % methylation as the interaction between IR exposure and units of M.CviPI. This is likely due to the latter model having reduced power to detect differences than modelling % accessibility. Modeling using metilene (as % Gm$^5$C accessibility) or methylkit (as methylated/unmethylated counts) identified the fewest DARs. When regions were required to have at least 3 GC sites, we found that the binomial regression model and linear mixed modelling of % Gm$^5$C methylation produced similar results (**Supplemental Fig. 15C** and **15D**); increasing the minimum number of Gm$^5$C sites showed that the binomial regression either produced comparable results to or outperformed linear mixed models (**Supplemental Fig. 15C** and **15D**).

**Model selection for region-based analyses**

Based on the results of model testing for DMRs and DARs, it appeared that the binomial regression was the most appropriate test for region-based analyses. Linear mixed modelling may find the most significant DMRs and DARs, but it is limited as it identified mainly small regions of 2 or 3 sites, its response variable (% m$^5$C methylation or % Gm$^5$C accessibility) does not take into considering variability in coverage, and linear modelling also allows for predicted values outside the possible range (i.e. <0% and >100%). methylSig is unable to be used for GC accessibility analysis as it does not use covariates, and methylKit and metilene give inconsistent results between DMRs and DARs: methylKit finds many DMRs but very few DARs, while metilene generally underperforms and finds several hundred DARs but fewer DMRs (**Supplemental Fig. 14** and **15**). The binomial regression uses the coverage information rather than an estimate of % methylation or % accessibility, takes into consideration the differences in coverage by modeling each position within a region with a random intercept, and performs consistently for DMRs and DARs. We therefore set the following criteria for calling DMRs and DARs:

(1) Regions must contain at least 5 methylation sites, grouped together as illustrated in **Supplemental Fig. 6**;

(2) A binomial regression FDR-corrected $P < 0.05$, as calculated using SAS's GLIMMIX procedure; and

(3) An average difference in methylation or accessibility of 10% or greater.

We considered a higher minimum accessibility difference for DARs however most DARs detected using the binomial regression had average differences in accessibility of 20% or greater. In addition, as most significant DMRs had averages methylation differences between 10% and 20% we chose to use a 10% accessibility difference for consistency with the methylation analysis.

**Supplemental Table 9.** Summary of analysis algorithms tested

| Model | Input | Software implementation | Allows covariates? | Application |
|---|---|---|---|---|
| Credible methylation or accessibility difference | % methylation / % accessibility | n/a | n/a | DMC, DAC |
| Wald test, beta-binomial distribution | Read counts | DSS (Park and Wu 2016) | Yes | DMC, DAC |
| Fisher's exact test / Binary logistic regression | Read counts | methylKit (Akalin et al. 2012) | No (FET) / Yes (regression) | DMC, DAC, DMR, DAR |
| Beta-binomial regression | Read counts | methylSig (Park et al. 2014) | No | DMC, DMR |
| Binomial logistic regression | Read counts | methylSig (Park et al. 2014) | No | DMC, DMR |
| Mann-Whitney U | % methylation / % accessibility | Metilene (Jühling et al. 2016) | No | DMC, DAC, DMR, DAR |
| Binomial logistic regression | Read counts | SAS v9.4, LOGISTIC and GLIMMIX procedures | Yes | DMC, DAC, DMR, DAR |
| Fisher's exact test | Read counts | SAS v9.4, FREQ procedure | No | DMC |
| Cochran-Mantel-Haenszel test | Read counts | SAS v9.4, FREQ procedure | Yes, as stratifying variables | DAC |
| Linear mixed model | % methylation / % accessibility | SAS v9.4, MIXED procedure | Yes | DMR, DAR |

DMC=differentially methylated cytosine; DAC=differentially accessible cytosine; DMR=differentially methylated region; DAR=differentially accessible region

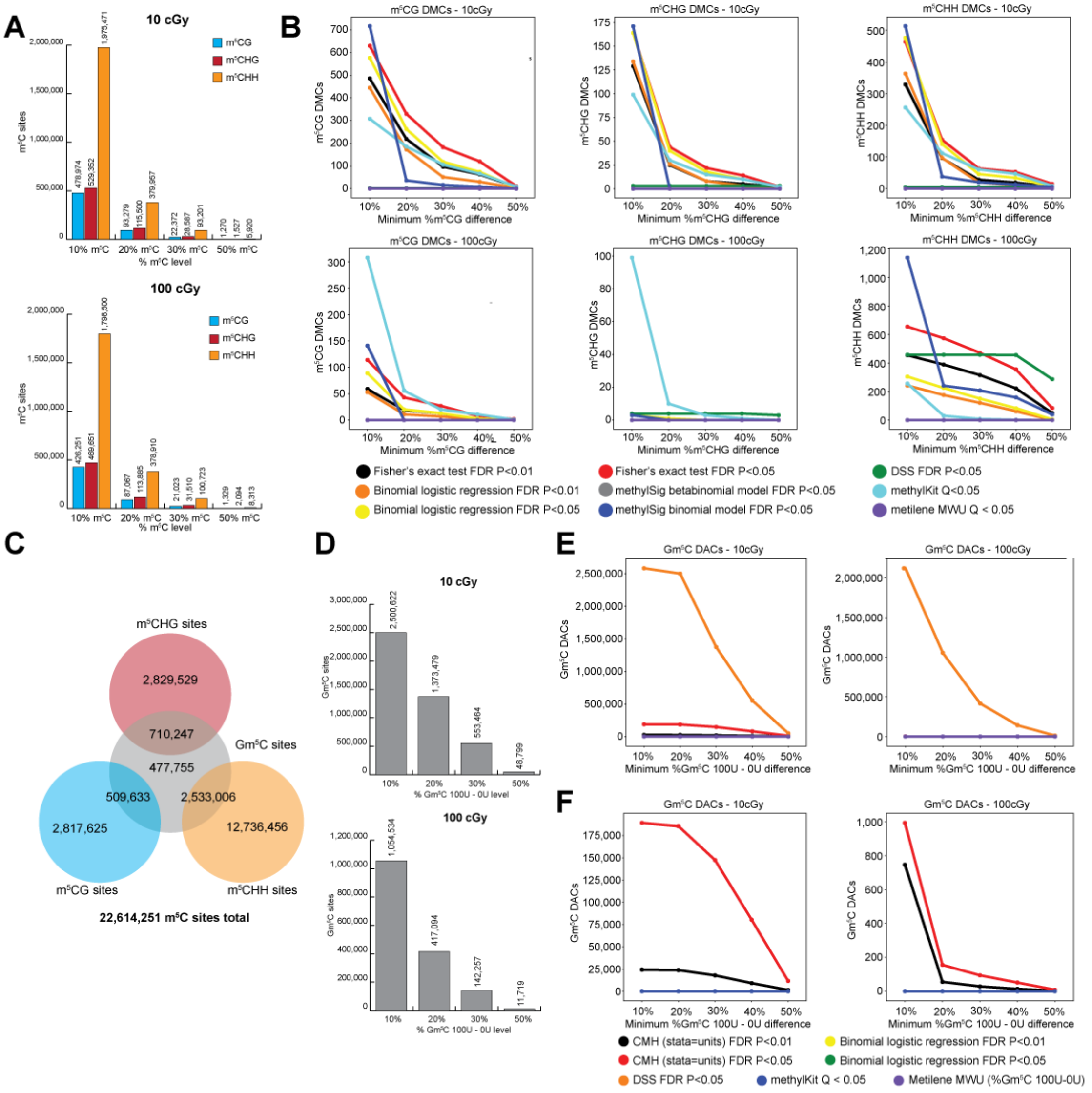

**Supplemental Figure 13.** Site-based tests for differential methylation and accessibility. (**A**) Distribution of number of endogenous m$^5$C sites by methylation (≥10%/≤-10%, ≥20%/≤-20%, ≥30%/≤-30%, ≥50%/≤-50%) for 10 cGy and 100 cGy IR exposure 72 h after exposure. (**B**) Endogenous differentially methylated cytosines (DMCs) by methylation type (m$^5$CG, m$^5$CHG, m$^5$CHH) and minimum difference in % m$^5$C for each statistical model tested. (**C**) Distribution of m$^5$C sites tested. (**D**) Distribution of number of Gm$^5$C sites by accessibility level (≥10%/≤-10%, ≥20%/≤-20%, ≥30%/≤-30%, ≥50%/≤-50% Gm$^5$C 100 U-0 U M.CviPI difference) for 10 cGy and

100 cGy IR exposure 72 h after exposure. (**E**) Gm$^5$C differentially accessible cytosines (DACs) by minimum difference in % m5C for each statistical model tested. (**F**) Panel E excluding the DSS model.

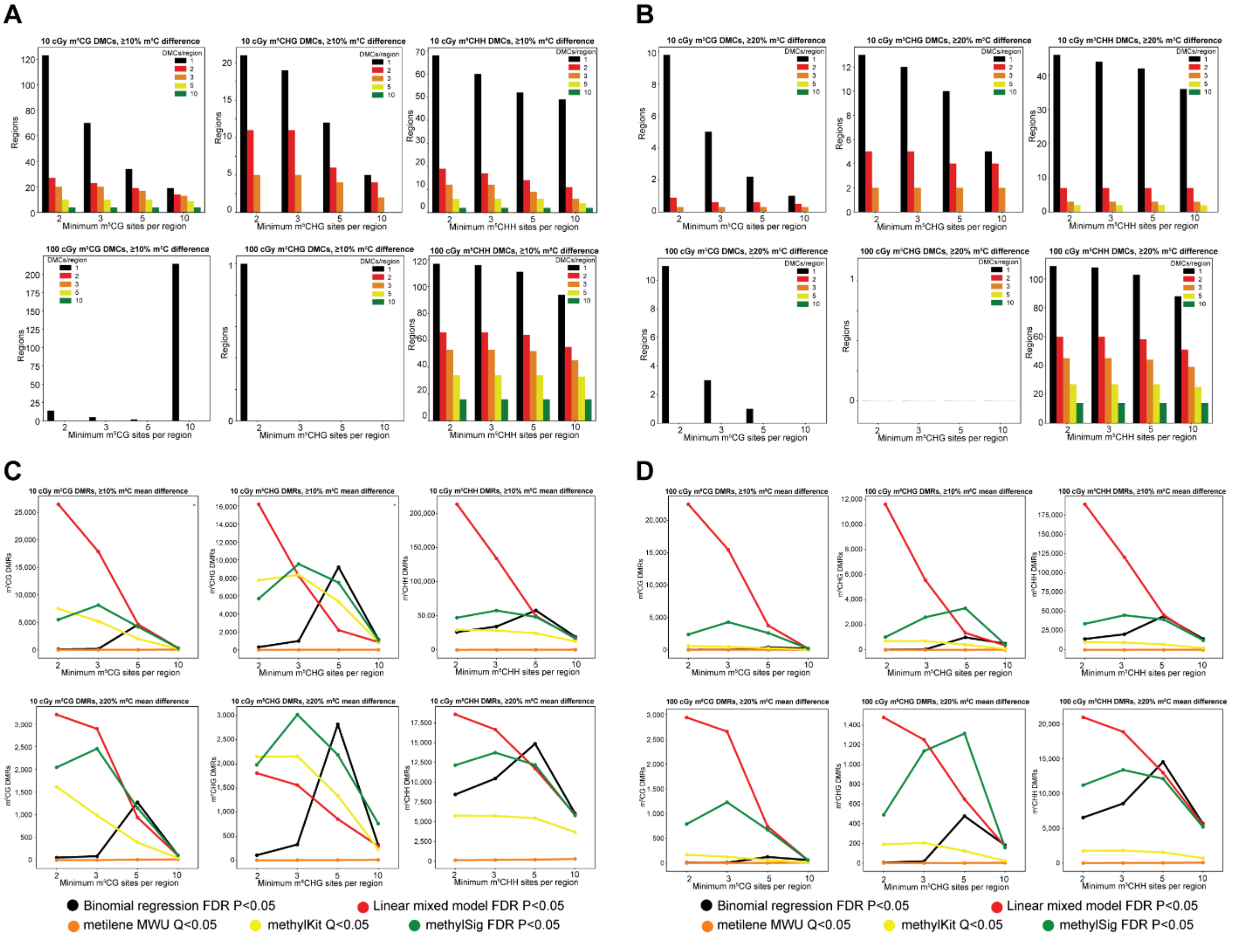

**Supplemental Figure 14.** Results of region-based differential endogenous methylation model testing. Distribution of regions of endogenous methylation by minimum m$^5$C sites per region and minimum number of DMC per region for DMCs with a minimum difference in %m$^5$C of (**A**) 10% or (**B**) 20%. Statistically significant differentially methylated regions (DMRs) by minimum m5C sites per region for each model test for DMRs with minimum average difference in %m5C of (**C**) 10% or (**D**) 20%.

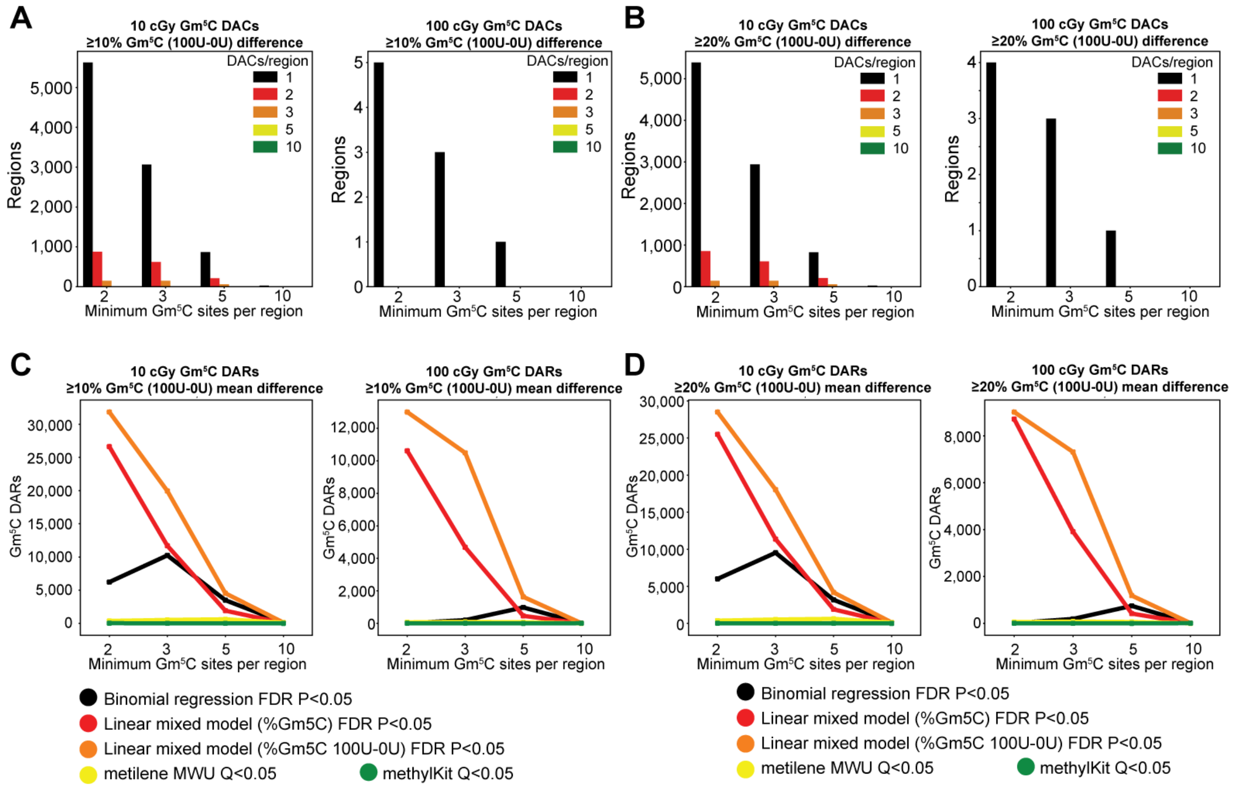

**Supplemental Figure 15.** Results of region-based differential accessibility model testing. Distribution of regions of Gm$^5$C methylation by minimum Gm$^5$C sites per region and minimum number of DAC per region for DACs with a minimum difference in %Gm$^5$C 100 U-0 U M.CviPI of (**A**) 10% or (**B**) 20%. Statistically significant differentially accessible regions (DARs) by minimum Gm$^5$C sites per region for each model test for DMRs with minimum average difference in %Gm$^5$C 100 U-0 U M.CviPI of (**C**) 10% or (**D**) 20%.
